# Prediction of single-cell chromatin compartments from single-cell chromosome structures by MaxComp

**DOI:** 10.1101/2024.07.02.600897

**Authors:** Yuxiang Zhan, Francesco Musella, Frank Alber

**Affiliations:** Department of Microbiology, Immunology, and Molecular Genetics, University of California Los Angeles, 520 Boyer Hall, Los Angeles, CA 90095; Institute of Quantitative and Computational Biosciences, University of California Los Angeles, Los Angeles, CA 90095; Department of Quantitative and Computational Biology, University of Southern California, 1050 Childs Way, Los Angeles, CA 90089, USA

## Abstract

The genome is partitioned into distinct chromatin compartments with at least two main classes, a transcriptionally active **A** and an inactive **B** compartment, corresponding mostly to the segregation of euchromatin and heterochromatin. Chromatin within the same compartment has a higher tendency to interact with itself than with regions in opposing compartments. **A/B** compartments are traditionally derived from ensemble Hi-C contact matrices through principal component analysis of their covariance matrices. However, defining compartments in single cells from single-cell Hi-C maps is non trivial due to sparsity of the data and the fact that homologous copies are typically not resolved. Here we present an unsupervised approach, named MaxComp, to determine single-cell **A/B** compartments from geometric considerations in 3D chromosome structures, either from multiplexed FISH imaging or from models derived from Hi-C data. By representing each single-cell structure as an undirected graph with edge-weights encoding structural information, the problem of predicting chromosome compartments can be transformed to an alternative form of the Max-cut problem, a semidefinite graph programming method (SPD) to determine an optimal division of a chromosome structure graph into two structural compartments. Our results show that compartment annotations from principal component analysis of ensemble Hi-C data can be perfectly reproduced as population averages of our single-cell compartment predictions. We therefore prove that compartment predictions can be achieved from geometric considerations alone using 3D coordinates of chromatin regions together with information about their nuclear microenvironment. Our results reveal substantial cell-to-cell heterogeneity of compartments in a cell population, which substantially differs between individual genomic regions. Moreover, by applying our approach to multiplexed FISH tracing experiments, our method sheds light on the relationship between single-cell compartment annotations and gene transcriptional activity in single cells. Overall our approach provides new insights into single-cell chromatin condensation, relationship between population and single-cell chromatin compartmentalization, the cell-to-cell variations of chromatin compartments and its impact on gene transcription.

**Author Summary:** Chromosome conformation capture and imaging techniques revealed the segregation of genomic chromatin into at least two functional compartments. Hi-C contact frequency matrices show checkerboard-like patterns indicating that chromatin regions are divided into at least two states, possibly a result of phase separation. Chromatin regions in the same state have preferential interactions with each other, often over extended sequence distances, while interactions to regions in the opposing state are minimized. Principal component analysis (PCA) on ensemble Hi-C contact frequency matrices can identify these compartment states. However, because the compartment annotations are derived from a cell population, this method cannot provide information about compartments in single cells. Here in this study, we introduce an unsupervised method to predict single-cell compartments using graph-based programming, which utilizes only structural information in single cells. Our results demonstrate that PCA-based ensemble compartment annotations can be reproduced as population averages of our single-cell compartment predictions. Moreover, our results reveal the cell-to-cell heterogeneity of compartments in a cell population, which shows significant disparities among different chromatin regions. Moreover, by applying our approach to multiplexed FISH tracing experiments, our method reveals the relationship between single-cell compartment annotations and gene transcriptional activity in single cells. Finally, our approach also allows us to relate chromatin structural features in single cells with compartment properties. Comparison with other existing approaches showed that our method produces overall better compartmentalization scores in single cells.

## Introduction

The development of chromosome conformation caption [1–3] and imaging techniques [4–8] provides tools to increasingly understand local folding and organization of chromatin regions across different scales from nucleosomes to chromatin territories. Hi-C studies revealed the existence of chromatin loops and topological associating domains (TADs) [9–13]. Moreover, chromatin regions segregate into at least two chromatin compartments, likely driven by protein factors that induce phase separation [14–16]. A widely used method to determine chromatin compartmentalization is to apply principal component analysis to the correlation matrix of an ensemble Hi-C contact frequency matrix [1,2]. The first principal eigenvector is used to assign chromatin regions with positive eigenvalues to the **A** compartment, which correlate with open, transcriptionally active chromatin. Chromatin regions with negative eigenvalues are assigned to the **B** compartment, correlating with closed, transcriptionally inactive heterochromatin. **A**/**B** chromosome compartment profiles correlate well with other experimental signals of transcriptionally active and repressive chromatin, including histone modifications from chip-seq data [17] and genomic information such as the GC content and CpG density [18].

However, most current analysis of chromatin compartments focused on ensemble datasets, describing chromatin properties averaged over a large population of cells. With the development of single-cell genomics technologies such as single-cell Hi-C [19–22] and chromosome tracing techniques from multiplexed FISH imaging [4–8], it is becoming increasingly important to characterize chromatin compartments at the single-cell level. However, direct assignment of ensemble level annotations of compartments on single-cell structures is not appropriate due to structural variations at the single-cell level. Hence, an approach to determine compartments based on single-cell information is essential for better understanding structure and gene function relationship in single cells. Moreover, such an approach will reveal which chromatin regions are expected to show relatively high cell-to-cell variations of chromatin compartments.

To solve this problem, a few approaches have been developed to use either single-cell Hi-C matrices [22–24] or single-cell chromatin distance matrices [16,22] to infer **A**/**B** compartments in single cells. For instance, deep learning-based methods like scGHOST have been applied to learn from training datasets to define **A**/**B** compartments in single-cell Hi-C maps [24]. Other methods have used the amount of CG content of a chromatin region for single-cell compartment annotations [22]. However, these approaches either require high quality datasets or require ground truth labels for supervised learning.

Here we provide a strategy to determine single cell chromatin compartments based only on geometric considerations of chromosome structures, generated by modeling approaches [26] or by super-resolution multiplexed FISH imaging [7,8]. Our approach assumes that chromosomal regions within the same compartment are generally closer to each other than regions in opposing compartments. By abstracting a chromosome structure with a graph, we can represent each chromatin region as a node and connect it with other nodes with edge weights based on their geometric locations with respect to each other and similarity in structural features. Segregation of chromosomal regions into two compartments can then be modeled as the “Max-cut problem”, a graph theory-related problem, which optimizes a cut through a set of edges such that the total weights of the cut edges will be maximized [27,28]. We name this method MaxComp from heron.

The resulting single-cell compartment assignments not only conform with experimental compartment profiles obtained from ensemble Hi-C matrices, but also agree with considerations about structural properties and gene transcription activities from single-cell observations.We assessed our predicted single-cell compartments by comparing the frequency with which a chromatin region is assigned to the **A** or **B** compartment in the population of cells with the eigenvalue PC1 of the first principle component obtained from the traditional PCA analysis of ensemble Hi-C data. We observe a high correlation between these two profiles, which validates that single-cell compartment predictions by MaxComp are meaningful at the ensemble level. We also assessed single-cell compartment annotations by computing compartmentalization scores. MaxComp predictions show the highest compartmentalization scores and outperforms other methods, such as Hi-C-based PCA, distance-based PCA using average distance matrices, as well as a method using average CG content of a chromatin region’s spatial neighborhood for compartment prediction. Moreover, our single-cell compartment assignments agree with known properties of active and inactive chromatin, including preferences in nuclear positions, chromatin fiber condensation, and distances to nuclear bodies, which confirmed expected trends in known euchromatin and heterochromatin regions. Finally, we assessed single-cell compartment annotations with single-cell gene transcriptional data, which confirmed an increased transcriptional activity of genes when they were predicted in the **A** rather than in the **B** compartment in a single cell.

Our results emphasize that single-cell compartments can be determined by geometric considerations from single-cell chromosome structures. Moreover, our approach is robust and can be successfully applied to chromosome structures imaged at low resolution and coverage. Furthermore, our method proves that MaxComp can directly determine **A** and **B** compartment annotations without applying statistical inference, which would require certain prior knowledge or deep learning methods, where a large training dataset is needed.

Finally, our results indicate considerable cell-to-cell variation in compartment annotations for some chromatin regions, emphasizing the need to determine chromatin states in single cells rather than apply chromatin states derived from ensemble annotations.

## Results

### Formulation of compartmentalization problem by Max Cut

Our goal is to partition all chromatin regions in a single-cell chromosome structure into the active **A** and inactive **B** compartments based on single-cell geometric considerations without the knowledge of ensemble based **A**/**B** compartment annotations from Hi-C data. The assumption is that chromatin within the same compartment shows increased interaction propensity, thus overall smaller distances to each other, while interactions of chromatin in different compartments are less favorable. We also expect chromatin regions to be more likely surrounded by other regions of the same compartment in 3D space, leading to spatial segregation between **A** and **B** compartments. Furthermore, we expect regions of the **A** compartment to be located more frequently around nuclear speckles than regions of **B** compartment, as high transcriptional activity has been observed in the vicinity of nuclear speckles [29,30] (**Fig. 1A**).

**Figure 1:**
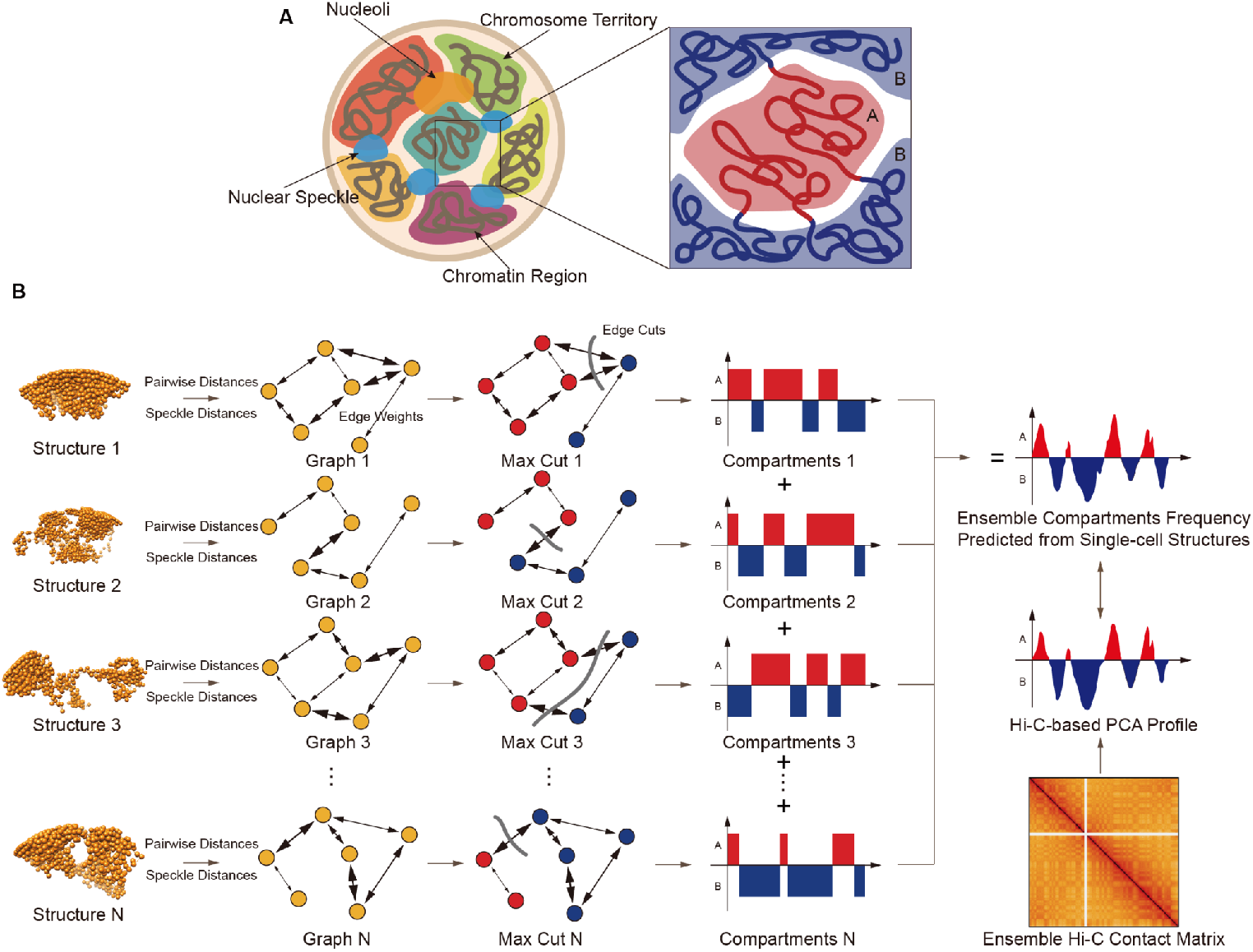
Overview of the MaxComp algorithm working on single-cell structures. **A**, The assumption of chromatin compartmentalization is based on two parts: Firstly, chromatin regions belong to the same compartment have higher contacts than those from the different compartment; Secondly, chromatin regions of compartment **A** are spatially closer to nuclear speckles than regions of compartment **B. B**, Every single-cell structure is transformed to an undirected graph which can be represented by an adjacency matrix whose edge weights are decided by its pairwise distances and speckle distances. Max Cut is then applied to the matrix to generate two partitions of nodes which are the prediction of the single-cell compartments of the structure. Ensemble compartment frequency can be calculated by combining a population of single-cell profiles.

To achieve our goal we represent each single-cell 3D chromosome structure (either from structure models or imaging experiments) as an undirected graph *G* = (*V, E*) with each chromatin region as a vertex *V* connected by edges with weights that encode information about their 3D distance and differences in their relative nuclear environment. Specifically, the edge weight between two genomic regions *i* and *j* is derived from their spatial distance normalized by the expected value based on their sequence separation as well as the z-score difference in their observed distances from the nearest nuclear speckle (Methods). Our goal is then to divide this graph into two subgraphs, which maximize the sum of their connecting edge weights, so that nodes within the same subgraph can be assigned to the same compartment and labeled as **A**/**B** according to the average speckle distance. This task can be achieved by solving a graph theory problem named the maximum-cut problem (Max-cut for short), which is NP-hard [31,32]. After solving the Max-cut problem for a given single-cell structure graph, we then assign **A/B** compartment annotations to each chromatin region. Each chromosome structure is then characterized by a single-cell compartment profile, delineating regions to either **A** or **B** compartment at the single-cell level.

### Relaxation and approximation of the Max-cut algorithm in MaxComp

We formulate the prediction of chromatin compartments in single cells as a Max-cut problem, which aims to find two subgraphs where the total weight of edges connecting them will be maximized. However, the Max-cut problem itself is NP-hard, and thus cannot be solved by a polynomial-time algorithm [31]. Besides greedy approaches, multiple approximation algorithms have been developed to reach a relatively high approximation ratio. Goemans and Williamson [27,28] have proven that certain relaxation and random projection techniques can increase the approximation ratio to about 0.878 to find a near optimal solution for each given Max-cut problem in polynomial time. The relaxation converts the original quadratic programming problem, where each node is represented by an indicator for its compartment type, into a vector programming problem where nodes are represented by vectors. This transformed problem can be reformulated as a semidefinite programming (SPD) problem, which aims to optimize a linear function subject to positive semidefinite constraints. Given the Laplacian matrix *L* which is a matrix representation of the graph *G* representing the target chromosome structure (Methods), the goal is to find a symmetric and positive semidefinite matrix ***A*** containing information of node labels with diagonal elements set to one so that we can maximize the objective function 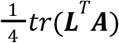 (Methods). We further demonstrate that maximizing intra-compartment similarity while simultaneously minimizing inter-compartment similarity can be accomplished using the same objective function, which proves that single-cell compartments are determined uniquely by ***L*** (Methods).

Ideally, we aim to obtain a strictly positive semidefinite matrix ***A***, enabling Cholesky decomposition to generate ***A*** = ***VV***^*T*^, where ***V*** contains row vectors representing nodes (i.e., genomic regions) in the hyperspace. However, generating a strictly semidefinite (SD) matrix requires lengthy computation times due to small convergence thresholds, especially on large graphs with hundreds of nodes. Therefore, we employ an approximation strategy to generate a close SD matrix with a larger convergence threshold. Subsequently, we apply lower-diagonal-lower (LDL) decomposition, a variant of Cholesky decomposition, which decomposes the target matrix into two triangular matrices and a diagonal matrix to obtained an approximated SD matrix ***A***′ = ***V***′ ***V***′^*T*^, facilitating the discovery of approximated row vectors (Methods). We prove that the difference in Euclidean norm between matrix ***V***′ containing approximated row vectors and matrix ***V*** is strictly governed by the difference in Euclidean norm between approximated matrix ***A***′ and matrix ***A***, ensuring minimal errors during approximation (Methods). Furthermore, Goemans and Williamson [27,28] introduced a random projection approach to iteratively generate hyperplanes, dividing all row vectors of ***V***′ into two groups labeled as compartment **A** and **B**. Using the adapted version of this algorithm, we are able to perform MaxComp several times faster, which is particularly suitable for high-resolution chromosome structures with several hundreds of chromatin regions.

### MaxComp prediction of single cell compartments from 3D chromosome structures

We apply our MaxComp approach to three different datasets. First, we use single-cell 3D genome structures generated by the integrative genome modeling (IGM) platform [26], using Hi-C [3], Lamin B1 DamID [33] and SPRITE [34] data as input information [26]. These structures are resolved at 200kb base pair resolution and also predict the distance of each chromatin region to the nearest nuclear speckle in each single cell, using a Markov clustering approach as described in [30]. Second, we apply MaxComp to 3D genome structures of human IRM90 cells from DNA multiplexed error-robust fluorescence in situ hybridization (MERFISH) experiments, a chromosome tracing method that images 3D genome structures for more than 7,000 single cells at a coverage of around 3Mb [7]. Thus 3D coordinates are available for chromatin regions separated by 3Mb in sequence distance across the entire genome. The method also imaged the locations of nuclear bodies, thus can provide information about an approximate distance of each chromatin region to nuclear bodies. Finally, we also apply our method to chromosome structures from DNA sequential fluorescence in situ hybridization (seqFISH+) imaging [8], which traces mouse embryonic stem cell (mESC) chromosomes at 1Mb coverage for 444 imaged cells (888 chromosome copies) and also provides relative locations of nuclear speckles within the same imaged cells.

### Applying MaxComp to chromosome structures from IGM genome structure modeling

First, we apply MaxComp on chromosome structures extracted from whole genome structures of H1-hESC cells generated by integrative modeling [26]. These structures also predict positions of nuclear bodies, such as nuclear speckles and nucleoli [26,30]. Specifically, we took 500 3D structures of chromosome 6 and 500 structures of chromosome 10 from single-cell H1-hESC whole genome structures. We then applied MaxComp to generate a single-cell compartment profile vector for each chromosome structure (Methods) (**Fig. 1B**). The input graph of a structure can be represented by an adjacency matrix, where the entry in row *i* and column *j* represents the weight of the edge connecting vertex *i* and vertex *j*. Adjacency matrices and predicted single-cell compartment profile vectors vary considerably across structures (**Fig. 2A and Fig 2B**, left column). Noticeably, along the chromosome sequence single-cell compartments show more frequent transitions between **A** and **B** compartments (**Fig. 2B**, left column), which can vary between individual cells, in comparison to the ensemble Hi-C derived compartment vector (**Fig. 2B**, right column). However, visualization of single-cell chromosome compartments in the 3D structures show a clear spatial segregation of **A**/**B** compartments, as indicated by extended clusters of chromatin regions in the same color (3D structures in **Fig. 2BC**, left panels).

**Figure 2:**
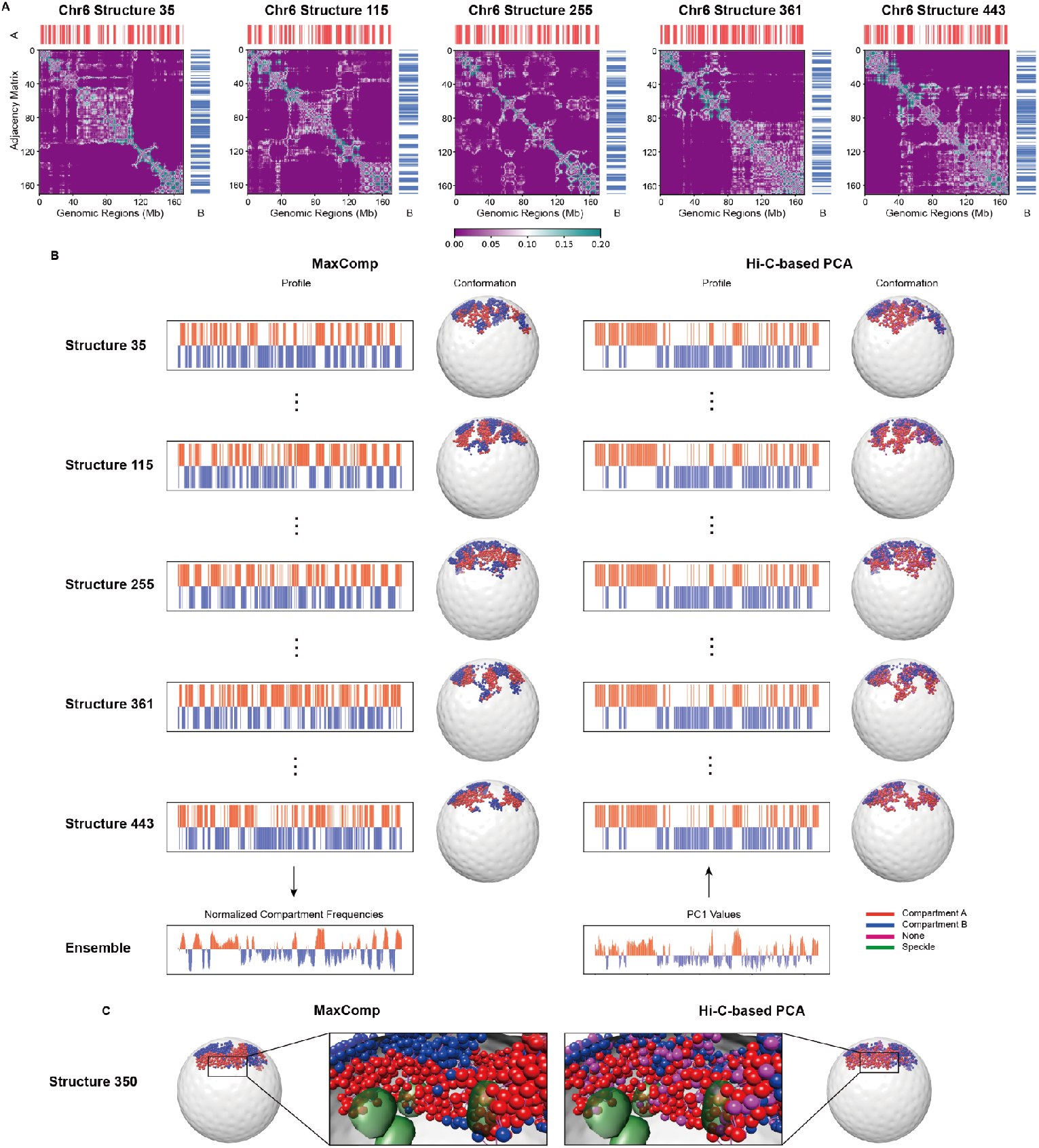
Selected example of model compartments predicted by MaxComp and the corresponding structures of H1-hESC Chr6. **A**, The predicted compartments for structure 35, 115, 255, 361, 443 of H1-hESC Chr6 together with the input adjacency matrices of the MaxComp approach. **B**, The compartment profile and 3D visualization of structure 35, 115, 255, 361, 443 of H1-hESC Chr6 colored by compartments (red in compartment **A** and blue in compartment **B**) from both experiment and the MaxComp prediction showed together with the nucleus envelope. **C**, Predicted speckles (green) showed together with the single-cell examples colored compartments labeled by MaxComp and the Hi-C-based PCA.

### MaxComp-predicted single-cell compartments reconstitute compartments from ensemble Hi-C

We first validate our MaxComp predictions by comparing the compartment frequencies (Methods) for each chromatin region with the PC1 values obtained from ensemble Hi-C matrices through principal component analysis (**Fig. 3A**). The absolute value of the compartment frequency of a given chromatin region measures how many times a chromatin region is assigned to either the **A** or **B** compartment across all single cells in the population (Methods). We observed high correlations (Pearson’s correlation >= 0.9) between the predicted normalized compartment frequencies and the PC1 principal component values for all studied chromosomes (**Fig. 3ABC**). Thus, the average chromosome compartment profiles over the population of cells reconstitutes the ensemble Hi-C derived compartment predictions, indicating that the single-cell predictions by MaxComp are meaningful at the ensemble level [1,3]. This agreement is an independent validation, since MaxComp only uses geometric considerations of 3D structures to derive the compartments (**Fig. 2B**).

**Figure 3:**
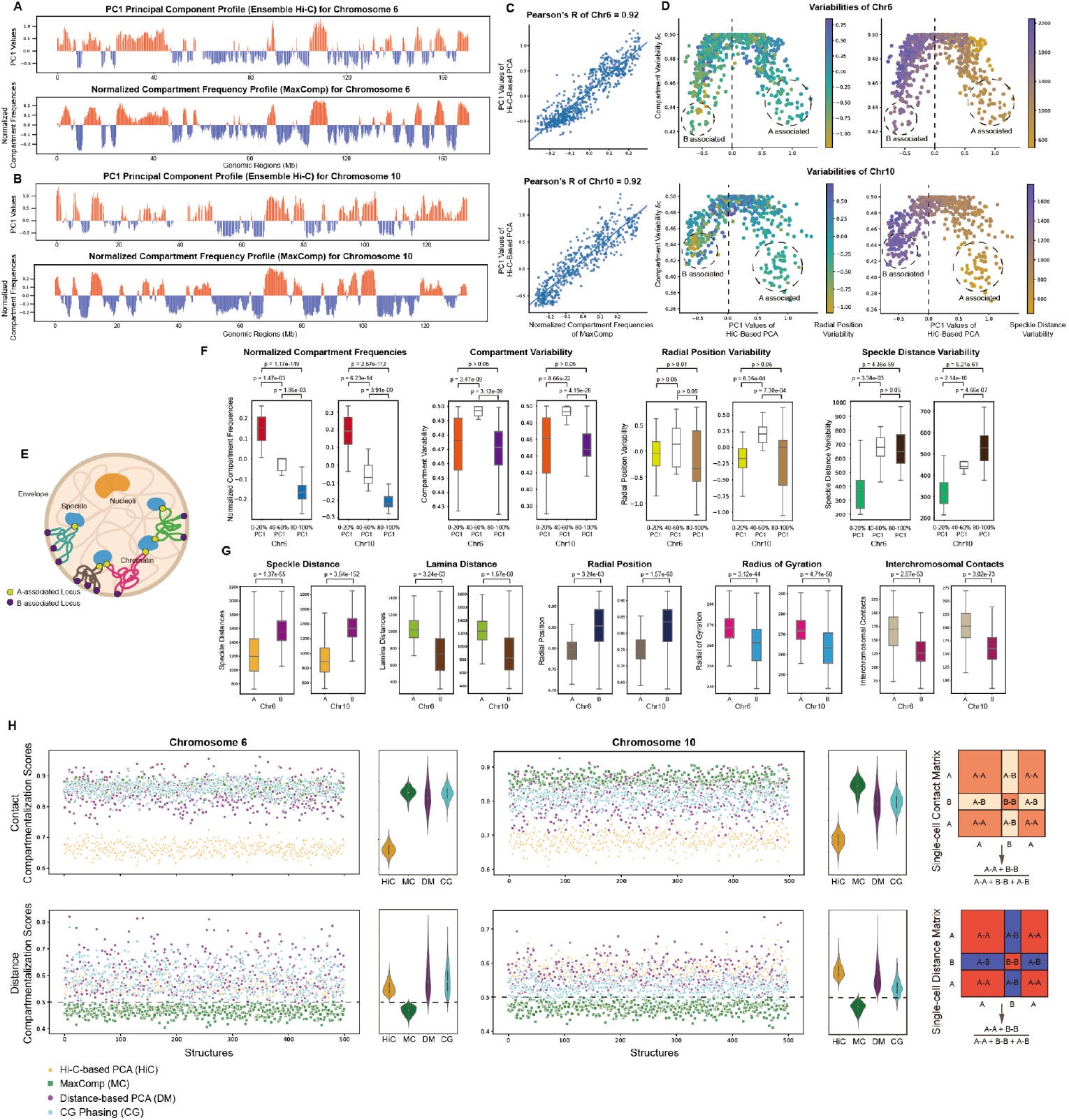
Prediction of model compartments by MaxComp and its comparison with other methods on H1-hESC Chr6 and Chr10. **A**, The experimental profile obtained from the Hi-C-based principal component analysis and the compartment profile predicted by MaxComp on 500 modeled structures of H1-hESC Chr6. **B**, The experimental profile obtained from the Hi-C-based principal component analysis and the compartment profile predicted by MaxComp on 500 modeled structures of H1-hESC Chr10. **C**, Scatter plot between the normalized compartment frequencies of MaxComp and the PC1 values together with the Pearson’s correlation coefficient between the two samples on each chromosome. **D**, Scatter plots of MaxComp single-cell compartment variabilities against PC1 values from the Hi-C-based PCA annotations for each chromosome. Each dot represents a genomic region colored by its value of cell-to-cell radial position variability in their radial positions (Left) or cell-to-cell speckle distance variability (Right). **E**, Illustration of **A**-associated locus and **B**-associated locus acting as anchors at speckles and envelope during chromatin folding. **F**, Comparison of compartment profile (p-value=8.84e-139 and 4.74e-140), compartment variability (p-value>0.05), radial position variability (p-value>0.01) and speckle distance variability (p-value=4.36e-59 and 5.21e-61) between top 20% PC1 locus and bottom 20% PC1 locus of H1-hESC Chr6 and Chr10. **G**, Comparison of speckle distances (p-value=1.37e-55 and 3.64e-152), lamina distances (p-value=3.24e-63 and 1.57e-60), radial position (p-value=3.24e-63 and 1.57e-60), radius of gyration (p-value=3.12e-44 and 4.71e-50) and interchromosomal contacts (p-value=2.87e-53 and 3.02e-73) between compartment **A** beads and compartment **B** beads on 500 structures of H1-hESC Chr6 and Chr10. **H**, Comparison of compartmentalization scores between the Hi-C-based PCA annotation, the MaxComp prediction, the distance-based PCA annotation and the CG phasing prediction for each structure from H1-hESC Chr6 and Chr10 showed together with the corresponding violin plots and illustration.

We observe that chromatin regions with intermediate Hi-C PC1 values (40 to 60 percentile; i.e., middle PC1 quintiles) show relatively low absolute compartment frequencies (close to 0) and thus show overall a significantly higher variability *δc*_*i*_ in their compartment assignments (Methods) (**Fig. 3D** left panel) than regions with large absolute PC1 values (p-value = 3.47e-09 and 3.12e-09 for Chr6 and p-value = 8.66e-22 and 4.19e-28 for Chr10 **Fig. 3F**). These regions often show significantly higher variability in their radial position in the nucleus than regions with high absolute PC1 values (p-value = 8.35e-04 and 7.30e-04 for Chr10, **Fig. 3D** radial position variability panel). These observations could indicate that these regions may have a higher transcriptional variability between single cells than regions with high absolute PC1 values, which tend to have the same dominant compartment assignments in the large majority of cells [30]. However, this is speculative at this point.

Therefore, regions with the highest and lowest PC1 value quintiles show the highest absolute compartment frequencies (**Fig. 3D**, left panels) and thus, the lowest compartment variabilities between cells than regions with intermediate PC1 value quintiles (p-value = 1.47e-03 and 1.86e-03 for Chr6 and p-value = 6.23e-14 and 3.91e-09 for Chr10, Compartment Variability panel in **Fig. 3F**) (**Fig. 3D**, left panels). These regions show also lower variability in their radial positions in the nucleus between cells and tend to be located either at the nuclear exterior lamina compartment or in the nuclear interior close to nuclear speckles, confirming previous observations that these regions can act as anchors for genome organization [30] (**Fig. 3D**) (radial position variability panel in **Fig. 3F**).

For instance, regions with highest normalized **A**-compartment frequencies show generally lower speckle distance variability and high affinity to nuclear speckles (**Fig. 3D**) (speckle distance variability panel in **Fig. 3F**).

Next, we test the robustness of the MaxComp pipeline with respect to the population size ranging from 10 to 500 single-cell chromosome structures. We found that 200 structures are sufficient to yield a Pearson’s correlation of at least 0.90 between MaxComp predicted normalized compartment frequency profiles and ensemble Hi-C compartment profiles (**Fig. S2ABC**). Even with only 10 structures the Pearrson’s correlation is already 0.64 (**Fig. S2ABC**). However, larger populations show the best performance and overall smoother normalized average compartment frequency profiles.

In summary, our analysis showed that single-cell compartment information can be determined from only 3D structural information and our method is robustly applicable to different chromosomes.

### Single-cell compartment predictions are consistent with expected structure properties

We further assess our compartment predictions by measuring several structural properties of chromatin to be predicted **A** and **B** compartments, including nuclear radial positions, chromatin fiber condensation, and distances to nuclear bodies.

Our analysis confirms several expected trends for euchromatic **A** and heterochromatin **B** compartment chromatin: First, we find chromatin predicted in **A** compartment to have significantly larger radius of gyration (p-value = 3.12e-44 for Chr6 and 4.71e-50 for Chr10), meaning that these regions are less condensed than **B** compartment chromatin (Radius of gyration panel in **Fig. 3G**). Also, compartment **A** regions have a significantly higher number of interchromosomal contacts in single cells than compartment **B** regions (p-value = 2.87e-53 for Chr6 and 3.02e-73 for Chr10). Thus, **A** compartment chromatin in single cells is more frequently located at the exterior of the chromosome territory (**Fig. 3G)**.

Also, chromatin in the **A** compartment are located more interior with smaller radial positions than those in the **B** compartment (p-value <= 1.57e-60 for Chr6,10) and subsequently smaller speckle distances (p-value <= 1.37e-55 for Chr6,10)(**Fig. 3G**).

All results are confirmed not only with structures of chromosome 6 and 10, but also other chromosomes, including chromosomes 8, 12, 15 and 18, which show very similar results (Pearson’s correlation >= 0.9 between predicted chromosome frequencies and Hi-C based PC1 profiles) (**Fig. S3ABC, S4ABC**).

### Single-cell compartments show high compartmentalization scores

To validate our results further, we calculate a compartmentalization score based on chromatin-chromatin contacts in single cell structures. The contact-compartmentalization score (CCS) is defined as the total number contacts between chromatin regions within the same compartments (intra contacts) divided by the number of all contacts irrespective of compartment assignments (Methods) (**Fig. 3H**). This value is averaged over all single-cell chromosome structures. A larger CCS score indicates a stronger segregation into the two spatial compartments, which is what we expect to be a more favorable state. We compare the contact compartmentalization scores from our MaxComp predictions with those from three other methods, namely the aforementioned ensemble Hi-C-based PCA analysis [2,3,8], a distance-based PCA analysis [4,25] and CG phasing [22] (Methods). The distance-based PCA method (DM) uses the average distance matrix calculated from all single-cell chromosome structures, and then applies principal component analysis on the average distance matrix to define **A**/**B** compartments. Similar to Hi-C-based PCA analysis, the resulting compartment annotations are then applied to all the chromosome structures. The CG phasing method determines for a given target region the total amount of CG DNA content from all chromatin regions located within its 3D spatial neighborhood to assign the A compartment to the target region.

MaxComp predictions achieve the overall highest average CCS compartment score among the three methods, especially in comparison to ensemble Hi-C-based PCA (HiC) and distance-based PCA (DM) compartment predictions (**Fig. 3H**). We also tested a compartment score based on distances instead of contacts between chromatin regions. The distance-compartmentalization score (DCS) is defined as the ratio of the average distances between chromatin regions within the same compartment and the average distances between all chromatin regions irrespective of their compartment annotation (Methods) (**Fig. 3H**). Here, a smaller DCS score indicates a more favorable spatial compartment segregation. Our results indicate that MaxComp compartment annotations produce a better spatial segregation between the two compartments at single-cell level as shown by the smallest DCS score in comparison to the other methods (**Fig. 3H**).

### Applying MaxComp to multiplexed FISH imaging datasets indicates the relationship between single-cell compartments and transcription signals

Next, we test MaxComp on chromosome structures imaged by integrated multiplexed FISH experiments [7,8]. These chromosome tracing experiments provide chromosome structures at considerably sparser coverage. For instance, chromosomes of human IMR90 cells are imaged at 3Mb step size in DNA MERFISH [7], while chromosomes in mESC cells are imaged at 1Mb step size in DNAseqFISH+ [8]. By combining multiplexed FISH chromosome tracing with immunofluorescence imaging these methods can also detect the locations of nuclear speckles and nucleoli in the same cells. Moreover, the datasets are also integrated with RNA-MERFISH and RNAseqFISH+, providing information about active transcription of specific genes in the same imaged cells [7,8].

We tested our method on 7,000 structures of chromosome 6 and chromosome 10 for IMR90 cells by DNA MERFISH [7]. Chromosome 6 is represented by a total of 55 imaged loci. Despite the relatively sparse coverage, MaxComp shows a good performance in predicting **A**/**B** compartment annotations. For instance, the averaged single-cell normalized compartment frequency profiles predicted by MaxComp have high correlations with the ensemble Hi-C-based PCA compartment profiles (Pearson’s correlation >= 0.8) (**Fig. 4ABC**). We also observe lower compartment variability for chromatin regions with high or low PC1 values, derived from independent Hi-C data analysis [2,3,8] (**Fig. 4D**). Also, **A** compartment chromatin shows significantly smaller speckle distances (p-value = 0.0 for Chr6 and 0.0 for Chr10) and larger lamina distances (p-value = 1.58e-102 for Chr6 and 2.20e-119 for Chr10) than compartment **B** chromatin (**Fig. 4E**).

**Figure 4:**
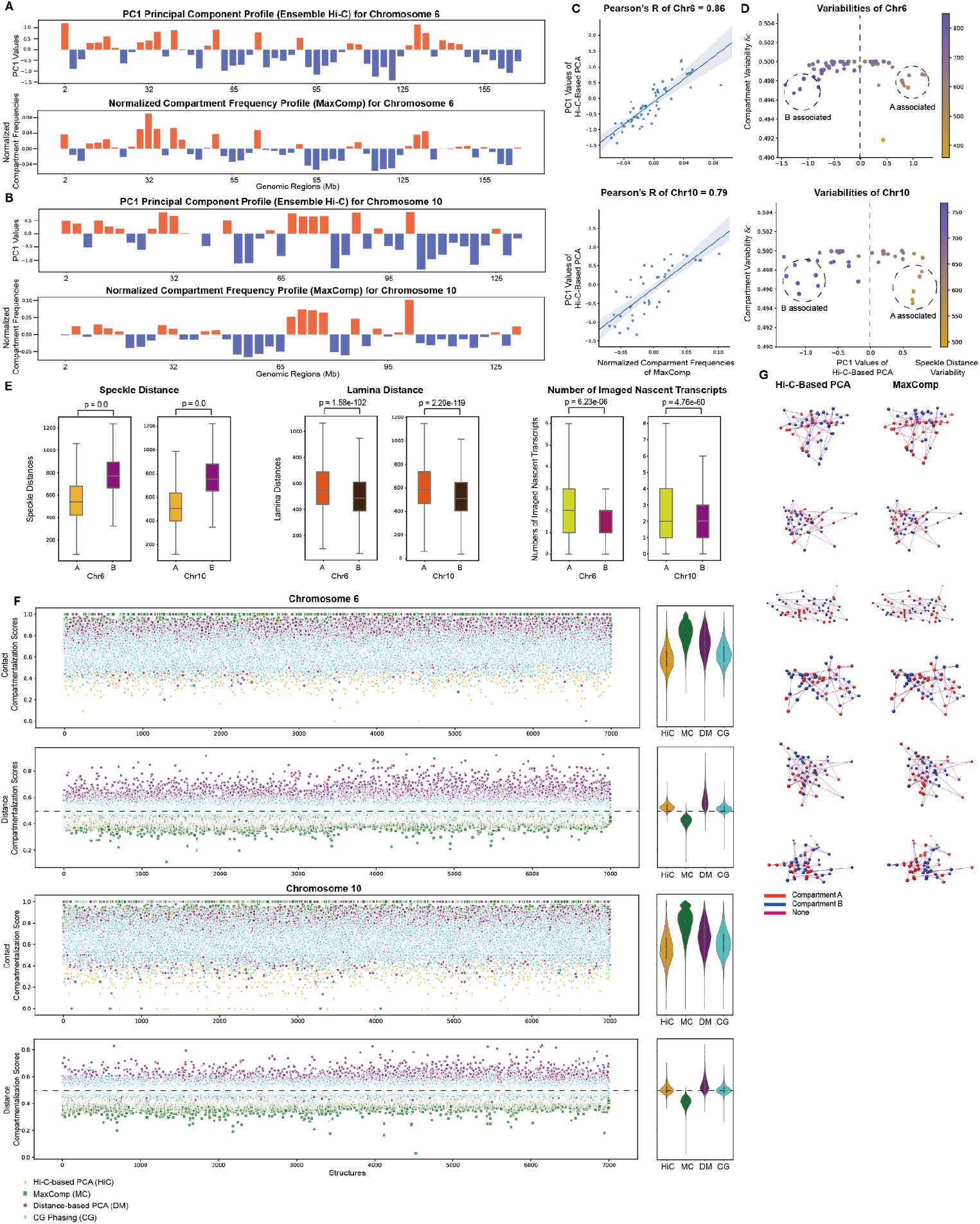
Prediction of DNA-MERFISH compartments by MaxComp and its comparison with other methods on IMR90 Chr6 and Chr10. **A**, The experimental profile obtained from the Hi-C-based principal component analysis and the compartment profile predicted by MaxComp on 7,000 DNA-MERFISH structures [7] of IMR90 Chr6. **B**, The experimental profile obtained from the Hi-C-based principal component analysis and the compartment profile predicted by MaxComp on 7,000 DNA-MERFISH structures of IMR90 Chr10. **C**, Scatter plot between the normalized compartment frequencies of MaxComp and the PC1 values together with the Pearson’s correlation coefficient between the two samples on each chromosome. **D**, Scatter plot of compartment variabilities against PC1 values from the Hi-C-based PCA annotations on each chromosome. Each dot represents a genomic region colored by its value of speckle distance variability. **E**, Comparison of speckle distances (p-value = 0.0 and 0.0), lamina distances (p-value = 1.58e-102 and 2.20e-119) and numbers of imaged nascent transcripts (p-value = 6.23e-06 and 4.76e-60) between compartment **A** beads and compartment **B** beads on 7,000 DNA-MERFISH structures of IMR90 Chr6 and Chr10. **F**, Comparison of compartmentalization scores between the Hi-C-based PCA annotation, the MaxComp prediction, the distance-based PCA annotation and the CG phasing prediction for each structure on IMR90 Chr6 and Chr10 showed together with the corresponding violin plots. **G**, The 3D visualization of 6 selected DNA-MERFISH structures of IMR90 Chr6 colored by compartments (red in compartment A and blue in compartment B) from both ensemble PC1 and Max-cut prediction.

A benefit of chromosome tracing by DNA MERFISH is that nascent gene transcription can be measured concurrently for a selected group of genes in the same cell by RNA MERFISH imaging [7]. Interestingly, we find for all tested genes a higher nascent transcription signal (i.e., number of imaged nascent transcripts in a cell) in those structures where the gene is predicted to be in the **A** compartment in comparison to cells where the same gene locus is predicted to be in the **B** compartment (p-value = 6.23e-06 for Chr6 and 4.76e-60 for Chr10) (**Fig. 4E**). Overall genes in the **A** compartment are more likely associated with active transcription, supporting studies about the role of nuclear compartmentalization for gene transcription [35]. However, our results also indicate that transcription can also occur for genes in the **B** compartments, confirming recent studies using RD-SPRITE, which simultaneously maps the 3D genome structure and nascent RNA transcription genome-wide [36] (**Fig. S6A**). Nevertheless, the average transcription profile (i.e. number of imaged transcription spots averaged across the population of cells) shows good correlation with the predicted **A**/**B** compartment profile (**Fig. S6B**). Similar results are also found in other chromosomes, such as chromosome 10 (**Fig. S7AB**).

Finally, we find that both compartmentalization scores, CCS and DCS, are significantly better for compartments predicted by MaxComp than those predicted by the distance based PCA (DM), single-cell CG phasing method or ensemble based Hi-C PCA (Hi-C) (**Fig. 4F**), confirming our previous observations.

Next, we applied MaxComp to chromosome tracing data from DNA SeqFISH+ of the mESC cell line [8]. These structures were imaged genome-wide at 1Mb coverage. Also here, we found similar results with high Pearson’s correlations >= 0.8 between PCA-based compartment profiles from ensemble Hi-C and MaxComp predicted normalized compartment frequency profiles in both studied chromosomes. Also here, the CCS and DCS compartmentalization scores show good spatial segregation for the MaxComp compartments (**Fig. 5ABCDE**). The CG method shows similar performance to MaxComp, although with a lower contact-based compartmentalization score. However MaxComp performs substantially better when calculating the distance-based compartmentalization score (**Fig. 5E**).

**Figure 5:**
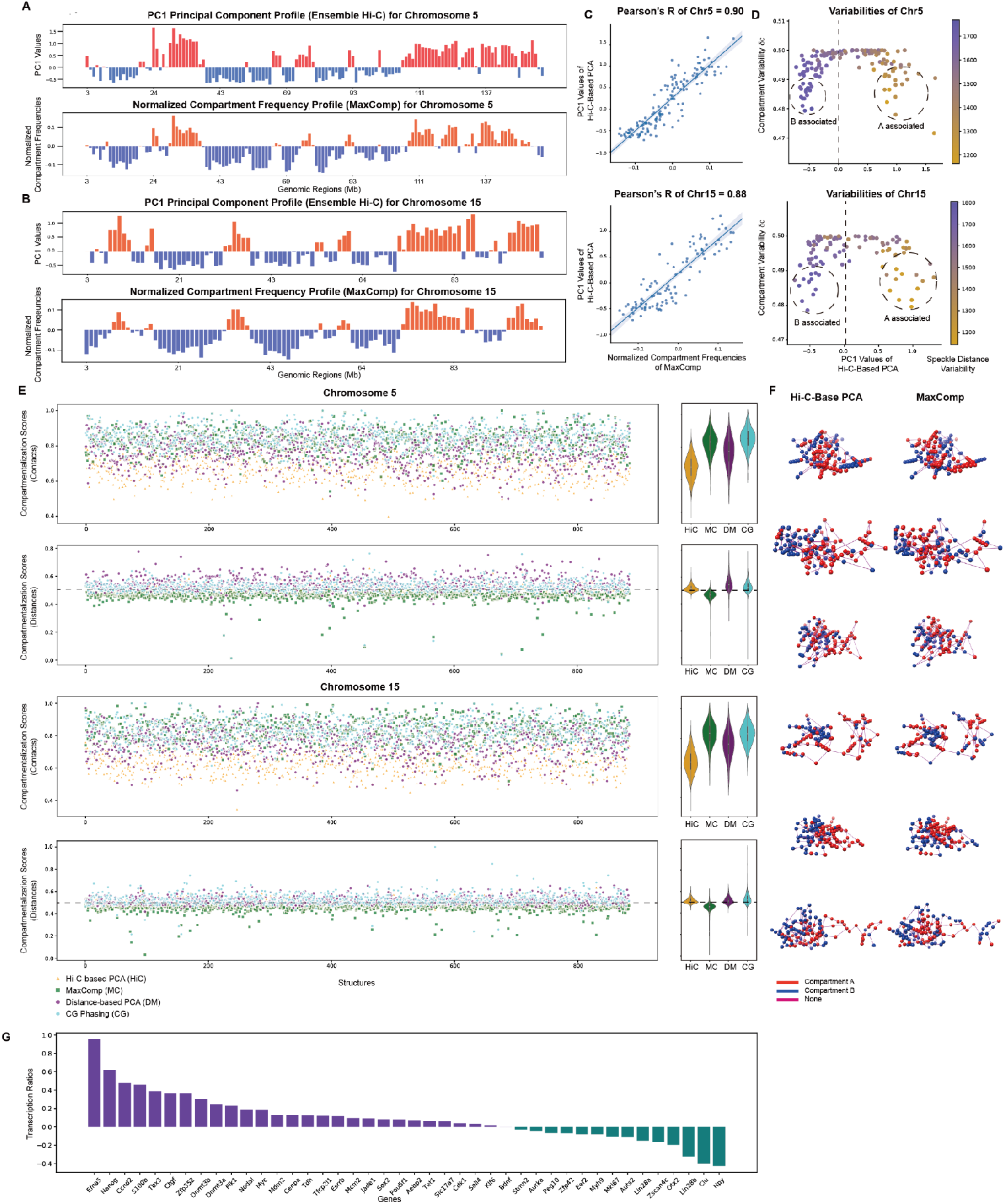
Prediction of SeqFISH+ compartments by MaxComp and its comparison with other methods on mESC Chr5 and Chr15. **A**, The experimental profile obtained from the Hi-C-based principal component analysis and the compartment profile predicted by MaxComp on 886 SeqFISH+ structures [8] of mESC Chr5. **B**, The experimental profile obtained from the Hi-C-based principal component analysis and the compartment profile predicted by MaxComp on 884 SeqFISH+ structures of mESC Chr15. **C**, Scatter plot between the normalized compartment frequencies of MaxComp and the PC1 values together with the Pearson’s correlation coefficient between the two samples on each chromosome. **D**, Scatter plot of compartment variabilities against PC1 values from the Hi-C-based PCA annotations on each chromosome. Each dot represents a genomic region colored by its value of speckle distance variability. **E**, Comparison of compartmentalization scores between the Hi-C-based PCA annotation, the MaxComp prediction, the distance-based PCA annotation and the CG phasing prediction for each structure on mESC Chr5 and Chr15 showed together with the corresponding violin plots. **F**, The 3D visualization of 6 selected SeqFISH+ structures of mESC Chr5 colored by compartments (red in compartment **A** and blue in compartment **B**) from both experiment and the MaxComp prediction. **G**, Log fold change of the average transcription level (number of mRNA transcript spots detected) of cells with both copies labeled with **A** (**A** cells) against the level of cells with both copies labeled with **B** (**B** cells) for each gene.

We also conducted analysis on 42 genes whose transcription levels have been determined in single cells [8]. To analyze these data we divide the structures in two groups: first, structures where the homologous gene copies are both predicted to be in the **A** compartment (**A** cells) and second, those structures where both gene copies were predicted to be in the **B** compartment (**B** cells). For each of these cells the nascent transcription level (number of mRNA transcript spots detected) for each of the 42 genes was measured by RNAseqFISH+ [8]. For each gene, we then calculate the transcription ratio as the average nascent transcription level for the genes in **A** cells over the average transcription level of the same gene in **B** cells (Methods). We find that 67% of all measured genes have positive transcription ratio, which shows that the same genes are expressed at higher levels in structures where the gene is predicted to be in compartment **A** (**A** cells) than in compartment **B** (**B** cells) (**Fig. 5G**). We find that the distributions of the normalized transcription levels of all 42 genes in **A** compartment are statistically significantly different from those in the **B** compartment (paired t-test p-value<0.05) (**Fig. S8**).

## Discussion

Experimental evidence from Hi-C and imaging data indicates that chromatin spatially segregates into at least two functional compartments. Compartment **A** corresponds to more open, transcriptionally active euchromatin and compartment **B** contains more condensed regions of heterochromatin. The spatial segregation of these compartments is detected in Hi-C contact frequency maps as a checkerboard like pattern indicating preferential interactions of chromatin with the same compartment and reduced interaction frequencies between chromatin in opposing compartments. Principal component analysis of ensemble Hi-C contact frequency maps allowed the division of chromatin into either **A** or **B** compartment, based on the PC1 eigenvalues of the chromatin region. However, this approach provides compartment annotations from chromatin properties averaged over a population of cells and therefore cannot reveal compartment information at the single-cell level. To better understand the underlying principles of chromatin compartment formation, it is important to detect compartments at single-cell level, and study if and how chromatin compartments vary between individual single cells.

To address this problem, we developed a graph-theory method called MaxComp, which determines chromatin compartments in single cells using only geometric considerations from 3D chromosome structures. By representing chromosome structures as an undirected graph, we show that MaxComp can optimally divide the structure graph into two subgraphs, which reproduce properties of the **A/B** compartments from ensemble Hi-C. The resulting single-cell compartment annotations produce compartment frequency profiles highly similar to the experimental compartment profiles obtained from PCA analysis of ensemble Hi-C data. Moreover, structural features of chromatin regions in single cells, such as radial positions and chromatin condensation (i.e. radius of gyration of a chromatin region) show significant differences between chromatin of predicted **A** and **B** single-cell assignments and are consistent with differences expected for euchromatin and heterochromatin segregation.

We show that our method is robust with respect to the resolution of chromosome structures. It can be successfully applied to low-coverage chromosome structures from DNA MERFISH and DNAseqFISH+ chromosome tracing experiments. Combined with nascent mRNA imaging, we can show that most genes predicted in the **A** compartment exhibit overall higher transcriptional activity than the same genes in the **B** compartment.

Therefore, we show that single-cell compartment predictions from geometric considerations of 3D chromosome structures are sufficient to determine meaningful compartment annotations in single cells.

Our analysis shows that the cell-to-cell variability of single-cell compartment annotations strongly correlates with the PC1 values from ensemble Hi-C compartment predictions. Chromatin regions with high absolute PC1 values show low compartment fluctuations. Instead, chromatin regions with intermediate PC1 values (i.e., small absolute values) show the highest fluctuations in compartment annotations between individual cells. Interestingly, these are also chromatin regions that show relatively high variability in their radial positions between individual cells. However, the high compartment fluctuations of these specific chromatin regions may also indicate a limitation of the two state compartment definition and that these regions may not be functionally clustered into either **A** or **B** compartments.

Moreover, our single-cell compartment annotations show good performance against other prediction methods, leading to more favorable compartmentalization scores and thus higher spatial segregation of **A** and **B** compartment chromatin.

Overall, our study emphasizes the importance of defining compartments at single-cell level, which can be achieved by using only geometric considerations in 3D chromosome structures. These structures can be derived from structure modeling or chromosome tracing experiments.

Our method is an unsupervised approach using “de novo” properties and does not rely on pre-trained ground truth labels, which are unknown. It can be applied broadly on chromosome structures at different coverage and resolutions. However, in future higher-resolution structures from imaging are needed to further assess the quality of compartment predictions.

## Materials and Methods

### Definition of Max Cut

Given an undirected graph *G* = (*V, E*), a cut in *G* is a subset *S*⊆*V*. Let 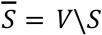, then the Max-cut problem is finding the cut *S* such that 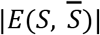 is maximized. Since the Max-cut problem is NP-hard, we cannot find a polynomial-time algorithm to solve the problem. However, multiple approximation algorithms which achieve approximate results exist, including greedy algorithms and local search. If we define the approximation ratio of algorithm *A* for the Max-cut problem by:

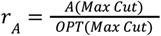

where *A*(*Max Cut*) is the result of algorithm *A* and *OPT*(*Max cut*) is the optimal result of the problem. Sahni and Gonzalez [31] comes up with an approximation algorithm that can achieve *r* _*A*_ = 0. 5, and many other algorithms which slightly increase the approximation ratio are then developed [28]. However, previous studies also have proved that the approximation ratio can be significantly improved if we regard the original problem as a quadratic programming problem [27,28].

### Quadratic programming problem

The relaxation and reformulation of the problem is from Goemans and Williamson [27,28]. Since finding a cut of a graph is equivalent to dividing the nodes of the graph into two group, we may use an indicator *x*_*i*_∈{1, − 1} to represent the group of node *i*. The objective of this problem is to find a division such that the sum of weights of all edges each connecting two nodes belonging to different groups is maximized. Then the Max-cut problem can be expressed as a form of quadratic programming problem:

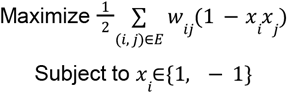

After solving out the indicators, we can classify all nodes that have the same indicator to the same group. Then we can define *S* = {*i*: *x*_*i*_ = 1} and 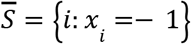

### Vector programming problem

A common approach to solve a hard problem is to relax some of the constraints and create a new relaxed problem. Thus, instead of representing every node by a one-dimensional indicator,we may relax the label of node *i* as a vector *v*_*i*_ ∈ *R*^*n*^ Then the Max-Cut problem can be relaxed as:

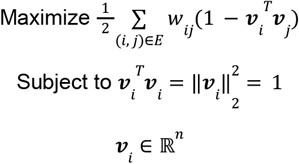

### Semidefinite programming problem

As we can observe in the vector programming problem, every vector *v*_*i*_ ∈ *R*^*n*^ actually presents in dot product form 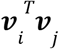 Hence, we may rewrite the dot product 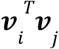 as *a*_*ij*_. Then the vector programming problem can be reformulated as:

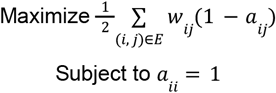

We may introduce a matrix ***A*** whose entries are *a*_*ij*_ Obviously ***A*** is symmetric according to the commutative properties of the dot product. Since 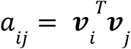 let ***V*** = (*v*_1_, *v*_2_, …, *v* _*n*_)^*T*^, then ***A*** = ***VV***^*T*^. For any nonzero vector *y* ∈ *R* ^*n*^, we also have:

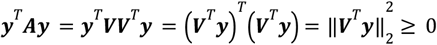

Thus, ***A*** is a symmetric and positive semidefinite matrix. We can reformulate the problem as a matrix programming problem:

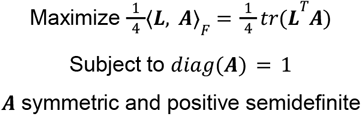

where ***L*** is the Laplacian matrix of the graph while ⟨***X, Y***⟩_*F*_ is the Frobenius inner product between matrix ***X*** and ***Y***. ***L*** can be directly calculated by ***L*** = ***E*** − ***W***, where ***W*** is the adjacency matrix and ***E*** the degree matrix of the graph.

### Formulation of the original problem

When predicting the compartments of chromosomes, we would like the beads in compartment A to contact more frequently with the beads in compartment **A** rather than the beads in compartment **B**. In other words, we want to maximize the contacts between the beads in the same compartments and minimize the contacts between the beads in different compartments. By transforming a single-cell structure to a graph, every node represents a bead and every edge represents the spatial connection between a pair of nodes. The weight between every two nodes is determined by their relative spatial information. According to the form of semidefinite programming problem, we want to maximize the sum of weights of edges connecting nodes from the different group, which is 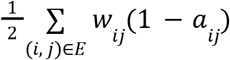, and minimize the sum of weights of edges connecting nodes from the same groups, which is 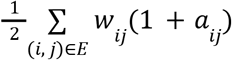. Then the original problem can be formulated as:

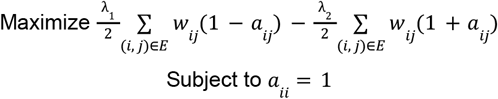

where *λ*_1_ and *λ*_2_ are factors of inter-compartment information and intra-compartment information with *λ*_1_ + *λ*_2_ = 1. The programming problem in matrix form can be formulated as:

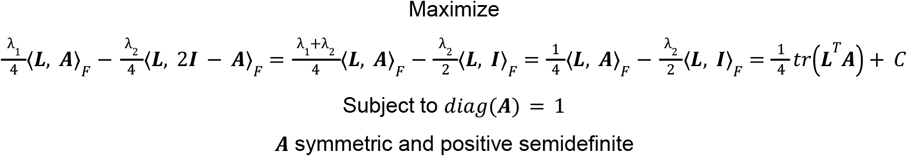

where ***I*** is the unit matrix. Since 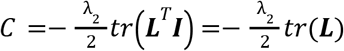 is a constant that doesn’t contain unknown variables, we may conclude that the objective function and the constraints of the original problem to find single-cell compartments is actually the same as the regular Max-cut problem. Hence, we have proved that considering both inter-compartment information and intra-compartment information is equivalent to considering only inter-compartment information. The implementation of the SDP version of the original problem is based on the python package cvxpy [37] and cvxopt https://github.com/cvxopt/cvxopt. We use the SCS solver [38] to solve the SDP problem and generate the resulting matrix ***A***.

### Decomposition by approximation and random projection

Since ***A*** = ***VV***^*T*^, after obtaining *A* by solving the above programming problem, we can further calculate ***V*** = (*v*_1_, *v*_2_, …, *v*_*n*_)^*T*^ by Cholesky decomposition. However, this decomposition can only be applied on a strictly positive semidefinite matrix. Due to the setting of convergence threshold, we may not be able to obtain a positive semidefinite matrix from the problem above. Instead, we will generate a symmetric matrix ***A*** which is very close to a positive semidefinite matrix. To avoid using Cholesky decomposition, we choose to apply LDL decomposition on ***A***:

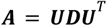

where ***U*** is an lower unit triangular matrix, ***D*** = *diag*(*d*_*i*_) is a diagonal matrix with diagonal entries *d*_*i*_ If ***A*** is an approximately positive semidefinite matrix, ***D*** may contain some negative entries with small values. To avoid negative values, we set 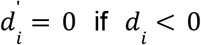 and 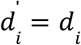 elsewhere to obtain a new diagonal matrix 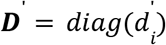 We define the square root matrix of ***D***′ by:

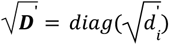

Since we know 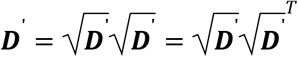 we can generate new matrix ***A***′by:

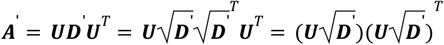

Hence, we can come up with a decomposition which is a variant of Cholesky decomposition by approximating diagonal entries of ***D*** and formulate the final vectors by:

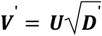

So that ***A***′ ***V***′ ***V***′^*T*^. Then the difference between ***A***′ and *A* can be evaluated by Euclidean norm of their difference and reverse triangular inequality:

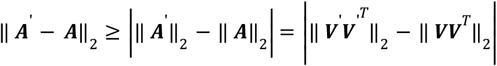

According to the property of Euclidean norm, we have 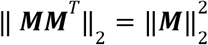 for any matrix ***M***:

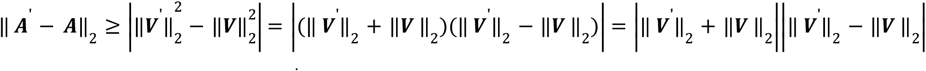

While the difference between ***V***′ and ***V*** is measured by Euclidean norm of their difference and triangular inequality:

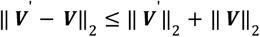

In all, we have:

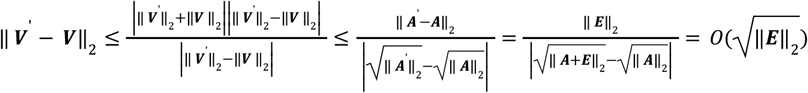

where ***E*** = ***A***′ − ***A*** donates the error between ***A***′ and ***A***. Hence, we have proven that if we can make the approximated matrix ***A***′ as close to matrix ***A*** as possible and reduce the error as much as possible, the resulting matrix ***V*** ^‘^ generated from decomposition will also be very close to matrix ***V***. In other words, we have proven ***A*** ≃ ***V***′***V***′^*T*^.

Let 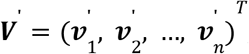, then for each node, 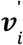 indicates its coordinates in the hyperspace. To classify the nodes into two groups, we can use a random hyperplane with normal ***r*** to cut the hyperspace into two sub-hyperspaces, where all vectors are divided into two groups with positive or negative dot products 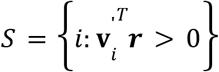 and 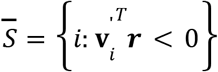 Using the two groups to form a vector **s** = (*s*_1_, *s*_2_, …, *s* _*N*_)^*T*^, where *s*_*i*_ = 1 if *i* ∈ *S* and *s*_*i*_ =− 1 if 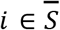, we may calculate the objective value of the Max-cut problem by:

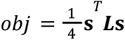

In order to obtain the best cut, we choose to apply random rounding multiple times until the objective value exceeds a threshold. Goemans and Williamson prove that the approximation ratio of random projection is about 0.878 which improves the performance significantly [27,28]. Hence, we may set the threshold to be 0. 878*OPT*(*Max cut*), where *OPT*(*Max cut*) indicates the optimal result.

### Transforming 3D structure to graph

The modeled population of H1-hESC genome structures is generated by integrative genome modeling (IGM) platform [26] with Hi-C [3], DamID [33] and SPRITE [34] as input, contains 1,000 whole-genome structures with coordinates of 30,332 200kb regions provided for each cell. Due to the large size of the graph and the long running time, we select 500 copies of each studied chromosome to perform the analysis.

The results of the compartments prediction depend on how we construct graphs from 3D structures by choosing proper edge weights. Firstly, the graph should be undirected. Secondly, since more contacts indicate closer distance between two beads, closer beads than expected should be assigned smaller weight between each other. Varoquaux et al [39] demonstrates that the expected spatial distance *d*_*ij*_ and genomic distance |*g*_*i*_ − *g*_*j*_ | between bead *i* and bead *j* at small ranges follows 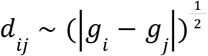. Since our analysis are chromosome-wide, given the 3D coordinates of bead *i* and bead *j* ***x***_*i*_ and ***x***_*j*_, we consider the ratio of observed distance against expected distance 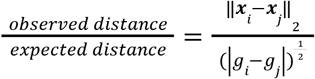 as part of the weight so that beads with longer distances than expected are more likely to be classified into different clusters.

We also use |*p* _*i*_ − *p* _*j*_ | as a scaler for each edge, where *p* _*i*_ and *p* _*j*_ is the distance of chromatin region *i* and *j* to the nearest speckle. The surface-surface between chromatin region and the nearest speckle is directly obtained from the models and the DNA-MERFISH datasets [7]. For the SeqFISH+ datasets [8] which provides only speckle density at certain loci, we estimate its speckle distance by assuming speckle density decays at quadratic rate:

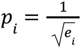

where *e*_*i*_ is the speckle density at loci *i*. In this way, edge weights are further normalized by their speckle affinity information and two beads are more likely to be classified into the same cluster if they have close speckle distances. The final weight between node *i* and node *j* can be formed by the product of the two normalized factors:

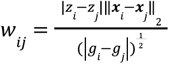

where ***x*** _*i*_ and ***x***_*j*_ are the 3D coordinates, *g*_*i*_ and *g* _*j*_ are the genomic positions, *z* _*i*_ and *z* _*j*_ are the z-scores of speckle distances *p* _*i*_ and *p* _*j*_ of bead *i* and bead *j*. Whatsmore, we choose to dropout edges with large weights, since compartments are more likely to be determined by local structural information. To dropout an edge, we set node *i* and node *j* not connected when ||***x***_*i*_ − ***x*** _*j*_||_2_ > 16*R*_*bead*_, where *R*_*bead*_ is the bead radius. For DNA-MERFISH and SeqFISH+ structures, we set *R*_*bead*_ = 100 *nm*. Finally, min-max normalization is applied to the weights so that all weights are located between 0 and 1:

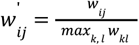

The matrix 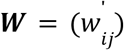 will be the final adjacency matrix of the graph, which is further used to generate the Laplacian matrix of the graph.

### Prediction of compartments

We apply the algorithm to every single graph and calculate the compartments for every single structure. Given two sets of nodes *S* and 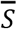 generated from the MaxComp algorithm, we set *S*_*a*_ = *S* if the average speckle distance of *S* is smaller than 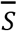, otherwise we set 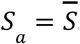

### Compartment profile vector

A compartment profile vector fo ***c*** _*k*_ = (*c* _*k*1_, *c* _*k*2_, …, *c* _*kN*_)^*T*^ for a given chromosome structure *k* is calculated based on the prediction results *S*_*a*_ and *S*_*b*_. We set *c* _*ki*_ = 1 if region *i* of structure *k* is predicted to be in compartment **A**, otherwise we set *c* _*ki*_ = 0.

### Ensemble compartment vector

We define the ensemble compartment vector as 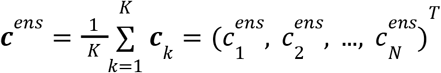 as the sum of all compartment profile vectors of all chromosome structures divided by the total number of structures. Each vector element 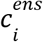 defines the fraction of times a genomic region *i* is assigned to the **A** compartment.

### Compartment frequency

The compartment frequency *f*_*i*_ of a genomic region *i* is defined as:

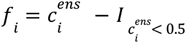

where 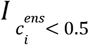 is an indicator function with 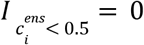 if 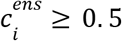 and 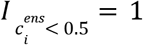 if 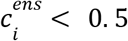. The absolute value of |*f*_*i*_| describes the fraction of times a genomic region *i* is predicted to be in the compartment of the majority of structures. For example, positive values indicate the compartment frequency of region *i* to be in compartment **A**, when the region is in compartment **A** in most of the structures. Negative values indicate the absolute value of compartment frequency of region *i* in compartment **B**, when the region is in the **B** compartment in the majority of structures. Subsequently, the compartment frequency profile vector for a chromosome is defined as:

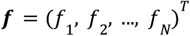

The compartment frequency vector can be compared with the PC1 profile vector calculated from the ensemble Hi-C experimental datasets.

### Normalized compartment frequency

The normalized compartment frequency 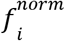 of a genomic region *i* is defined as:

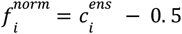

The normalized compartment frequency is used to calculate the correlation with PC1 values of ensemble Hi-C based compartment prediction.

### Compartment variability

The compartment variability *δc*_*i*_ for chromatin region *i* is calculated as the standard deviation of its compartment profile value across all *K* structures of the population:

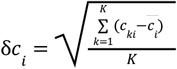

where *c*_*ki*_ is either 1 indicating that region *i* is in compartment **A** in structure *k* or 0 indicating that the same region *i* is in compartment **B**. 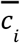 is the average value of *c*_*i*_ of region *i* across all *K* structures: 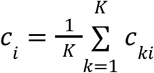. The larger the value of δ*c*_*i*_ is, the more variable are the compartment annotations of region *i* over the whole population.

## Structural features

### Radial position (RAD)

The radial position of a chromatin region *i* in structure *s* in a spherical nucleus is calculated as:

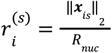

where ***x***_*is*_ is the the 3D coordinates of bead *i* in structure s, and *R*_*nuc*_ = 5 μ*m* is the nucleus radius. 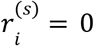 indicates the region *i* is at the nuclear center while 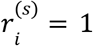 means it is at the nuclear surface. The radial position variability (δRAD) of region *i* in the population is calculated as:

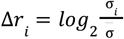

where σ_*i*_ is the standard deviation of the population of radial positions of region *i* and 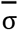 is the mean standard deviation calculated from all regions within the same chromosome of the target region.

### Radius of gyration (RG)

The local compaction of the chromatin fiber at the location of a given locus is estimated by the radius of gyration for a 1 Mb region centered at the locus. To estimate the values along an entire chromosome we use a sliding window approach over all chromatin regions in a chromosome. The radius of gyration for a 1 Mb region centered at locus *i* in structure *s*, is calculated as:

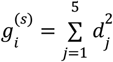

where *d*_*j*_ is the distance between the chromatin region *j* to the center of mass of the 1-Mb region.

### Distances to nuclear bodies

Various distances to nuclear bodies, including The speckle distance (SpD) and the lamina distance (LmD) for region *i*, are calculated by measuring the distance between the surface of each chromatin region *i* to the nearest speckle and lamina [30]. For each model, the locations of speckles are estimated by the geometric center of subgraphs related to the top 10% TSA-seq signal identified by Markov clustering [40] of the corresponding chromatin interaction network [30], while the locations of lamina is identified as the envelop. The SpD and LmD for DNA MERFISH is directly obtained from the datasets [7]. Similarly, the speckle variability (δSpD) of region *i* in the population is calculated as the standard deviation of the population of speckle distances of region *i*.

### Interchromosomal contacts

The calculation of inter-chromosomal contacts is similar to the calculation of contact frequency matrix but is based on a larger contact range. For a given 200kb region, its interchromosomal contacts is the total number of contacts with any target inter-chromosomal regions from the same genome structure within range *R*_*soft*_ = 1, 000 *nm*.

### Compartmentalization score

We define the compartmentalization score by the total contacts within the same compartments (intra-compartment contacts) divided by the total contacts of the whole structure (intra-compartment contacts + inter-compartment contacts). In other words, the score *s*_*c*_ contact-compartmentalization score (CCS) can be defined by for

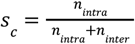

where *n*_*intra*_ indicates total unique intra-compartment contacts and *n*_*intra*_ indicates total unique inter-compartment contacts. Considering the differences in loci intervals, we define there is a contact when spatial distance between two beads are not larger than 3*R*_*bead*_ for model structures and SeqFISH+ coordinates [8] and 4*R*_*bead*_ for DNA-MERFISH coordinates [7]. By comparing different partitions of compartments (annotations from MaxComp or other methods), the ratio is able to tell us which partition is better. A larger score indicates larger intra-compartment contact percentage than inter-compartment one, which means the partition is more accurate and reasonable. Similarly we may assess pairwise distances instead of contacts to calculate the score *s*_*d*_ for distance-compartmentalization score (DCS) by

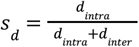

where *d*_*intra*_ is the sum of intra-compartment distances and *d*_*intra*_ indicates the sum of inter-compartment distances. A lower portion indicates smaller intra-compartment distances than inter-compartment ones, which contributes to better chromatin state segregation.

### Preprocessing of imaging tracing datasets

For the imaging dataset, 7,000 DNA-MERFISH copies of chromosome 6 and chromosome 10 from the IMR90 cell line are obtained from Su et al [7] together with their corresponding speckle distances, lamina distances, nucleoli distances and transcription profiles with transcription on or off (nascent transcript imaged or not) for genes measured by RNA MERFISH. All datasets are preprocessed by linear interpolation to remove missing values. Structural information of DNAseqFISH+ including coordinates, speckle densities and transcription information containing the number of detected spots corresponding to mRNA transcript for more than 40 genes from 444 cells (888 copies) of the mESC cell line are obtained from Takei et al [8]. We preprocess the datasets to generate reasonable speckle locations for each cell so that the experimental SON TSA-seq can be reconstructed in ensemble. Similarly, all datasets are preprocessed by linear interpolation to remove missing values. To avoid the impact of zero transcription, we remove the bottom 5% cells in the number of transcription spots for each gene when performing transcription ratio and paired change analysis. The genes are mapped to genomic regions nearest to their promoters in the reference genome [41]. For each gene, we first divide the cells into different groups, where **A** cells (**B** cells) indicate the corresponding locus on both copies is predicted to be in the compartment **A** (**B**). Then its transcription ratio *trans* is calculated by the log fold change of the average transcription level (number of mRNA transcript spots detected) in **A** cells *t*_*A*_ over the average transcription level in **B** cells *t*_*B*_ :

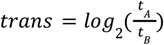

Normalized *t*_*A*_ and *t*_*B*_ are calculated in the same way for each gene after min-max normalization on the population-wide transcription levels to conduct paired comparison with paired t-test.

### Hi-C-based PCA

The Hi-C-based PCA profile ***c***_*exp*_ are obtained from the in-situ Hi-C dataset for H1-hESC (4DNESX75DD7R) [3], which are directly calculated by the largest principal component (the eigenvector corresponds to the largest eigenvalue) of the covariance matrix from the experimental ensemble Hi-C. For comparison, the PC1 values are mapped and averaged with regards to the nearest 200kb bins from the model. We measure the Pearson’s correlation coefficient *r*(***c, c***_*exp*_) between the non-zero values from the predicted normalized compartment frequency vector ***c*** and the experimental profile ***c***_*exp*_ for each studied chromosome. The Hi-C-based PCA profile ***c***_*exp*_ are obtained from Rao et al for IMR90 (4DNESSM1H92K) [2] and Takei et al [8] for mESC. Similarly, we use the nearest PC1 values for each imaged loci for comparison with predicted profiles.

### Distance-based PCA

Principal component analysis is mathematically performed by eigenvector decomposition on the input matrix, which can not only be applied on contact matrices, but also be adapted to pairwise distance analysis. The approach has been previously used by imaging related studies such as Wang et al [4] and Sawh et al [25]. We first normalize the mean distance matrix by pairwise genomic distances through fitted power-law function, and then calculate the pairwise Pearson’s correlation matrix between every row and column pair. Using it as the covariance matrix for PCA analysis, the resulting vector corresponding to the largest principal component with positive and negative entries can be used to generate compartments that are comparable with annotations from other methods.

### CG Phasing

Another frequently used approach is phasing by genomic information such as CpG density or CG content [22], where we have prior knowledge that high CG contents correspond to active compartments. We obtain CG contents for both hg38 and mm10 reference genomes from the UCSC genome browser [41]. For each loci, we calculate the mean CG contents from all locus (including itself) within its neighborhood (250 nm). The resulting vector measures the smoothed CG contents at single-cell levels, where the higher the value is the more likely the chromatin region belongs to **A** compartment. Eventually we may calculate the log fold change against the average to get **A**/**B** annotations by positive or negative signs as what we have explained in the MaxComp prediction.

## Acknowledgments

This work was supported by the National Institutes of Health (UM1HG011593 to F.A). We thank Lorenzo Boninsegna and Ye Wang for their help and useful discussions.

## Author contributions

Y.Z. and F.A. designed research. Y.Z. performed calculations and initial data analysis..Y.Z. and F.A. interpreted results and data analysis. F.M. and Y.Z. collected and processed imaging datasets. Y.Z. wrote software and documentation. Y.Z. and F.A. wrote the draft of the manuscript. Y.Z., F.A., F.M. edited the manuscript. All authors approved the final manuscript. F.A. secured funding.

## Figures

**Figure S1:**
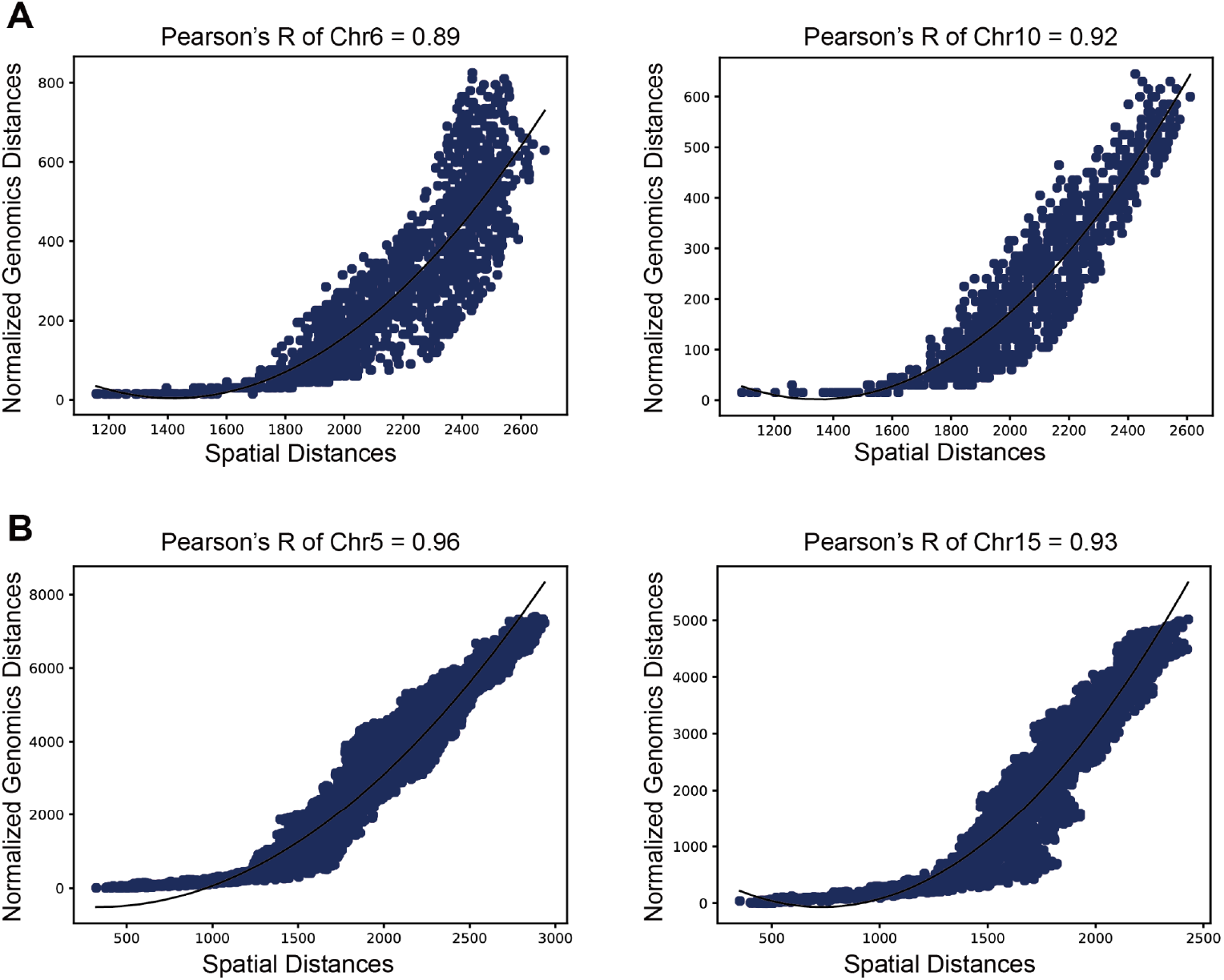
Correlations between average distances and normalized genomic distances. **A**, Scatter plots of average distances from DNA-MERFISH Chr6 and Chr10 [7] against the corresponding normalized genomic distances showed together with the fitted quadratic curves and the Pearson’s correlation coefficients. **B**, Scatter plots of average distances from SeqFISH+ Chr5 and Chr15 [8] against the corresponding normalized genomic distances showed together with the fitted quadratic curves and the Pearson’s correlation coefficients.

**Figure S2:**
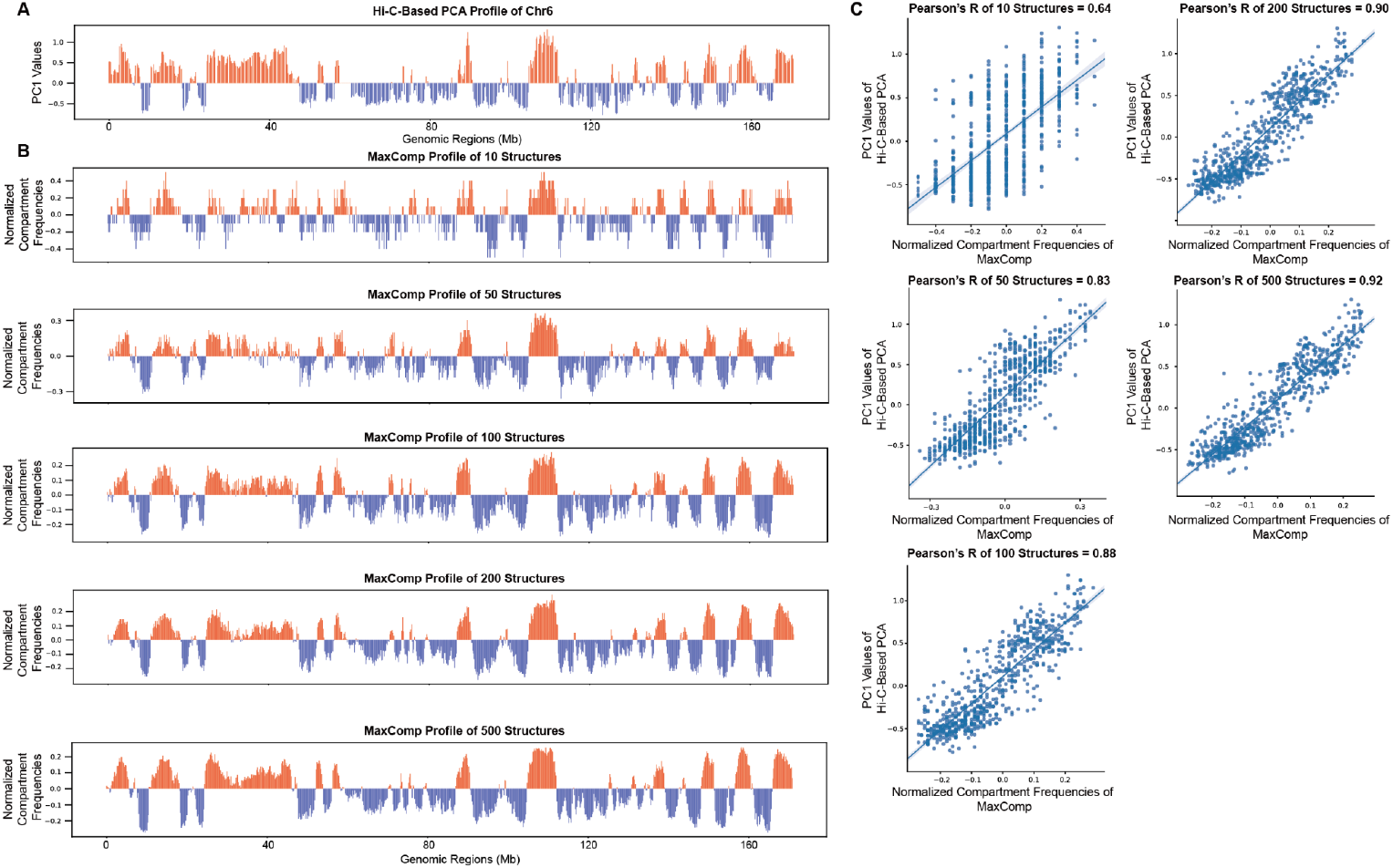
Comparison between population sizes on predicted profiles and their correlations. **A**, The experimental profile obtained from principal component analysis on ensemble Hi-C matrix. **B**, The predicted normalized compartment frequencies by MaxComp on populations of modeled Chr6 structures with different sizes. **C**, Scatter plots between PC1 values from the Hi-C-based PCA and various predicted compartment profiles showed with Pearson’s correlation coefficients. We observe increased value in the coefficient as population size grows larger.

**Figure S3:**
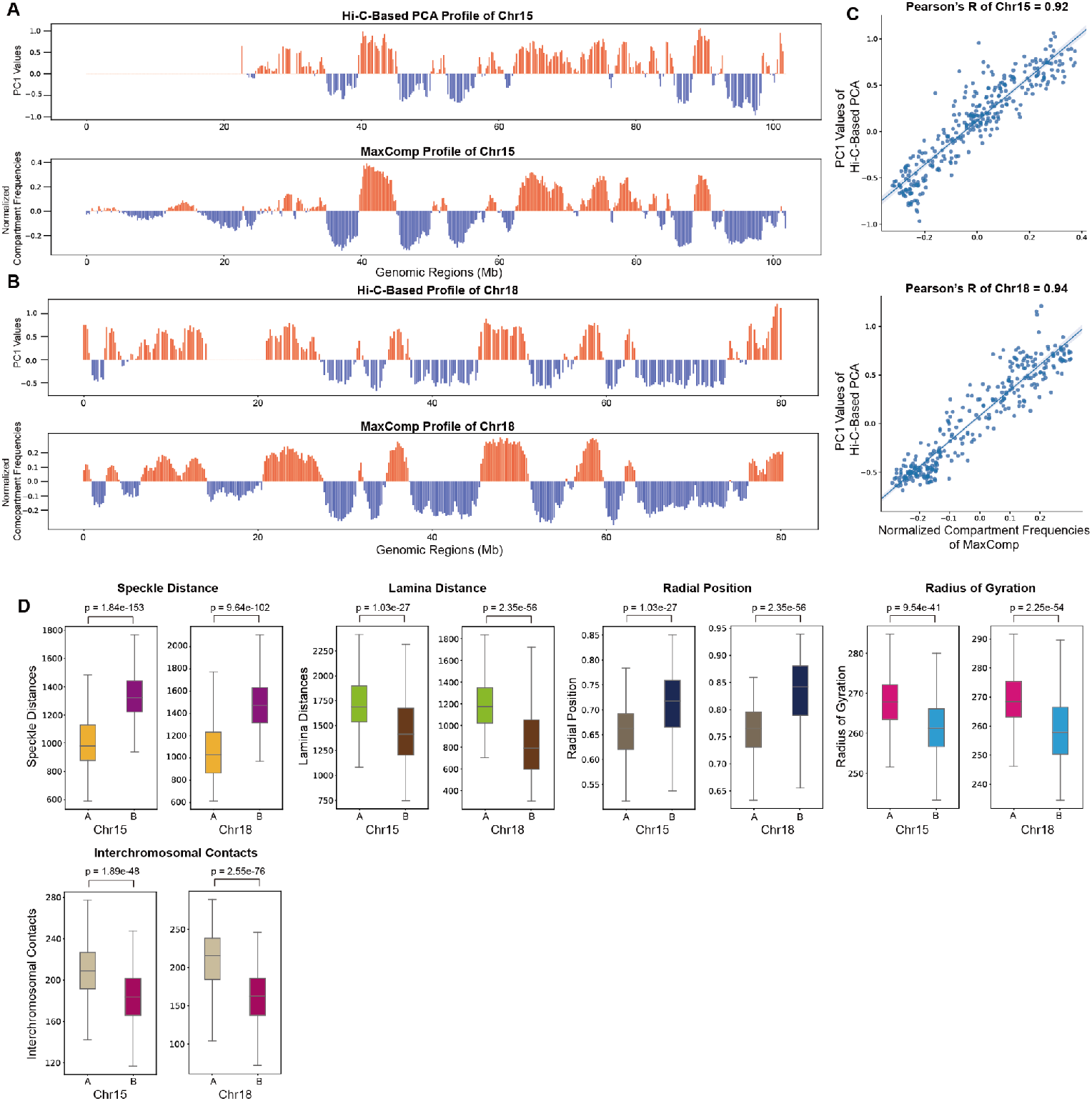
Prediction of model compartments by MaxComp and its comparison with the ground truth on H1-hESC Chr15 and Chr18. **A**, The experimental profile obtained from the Hi-C-based principal component analysis and the compartment profile predicted by MaxComp of 500 modeled structures of Chr15. **B**, The experimental profile obtained from the Hi-C-based principal component analysis and the compartment profile predicted by MaxComp of 500 modeled structures of Chr18. **C**, Scatter plot between the normalized compartment frequencies of MaxComp and the PC1 values showed together with the Pearson’s correlation coefficient between the two samples on each chromosome. **D**, Comparison of speckle distances (p-value=1.84e-153 and 9.64e-102), lamina distances (p-value=1.03e-27 and 2.35e-56), radial positions (p-value=1.03e-27 and 2.35e-56), radius of gyration (p-value=9.54e-41 and 2.25e-54) and interchromosomal contacts (p-value=1.89e-48 and 2.55e-76) between compartment **A** beads and compartment **B** beads on the population of structures of Chr15 and Chr18.

**Figure S4:**
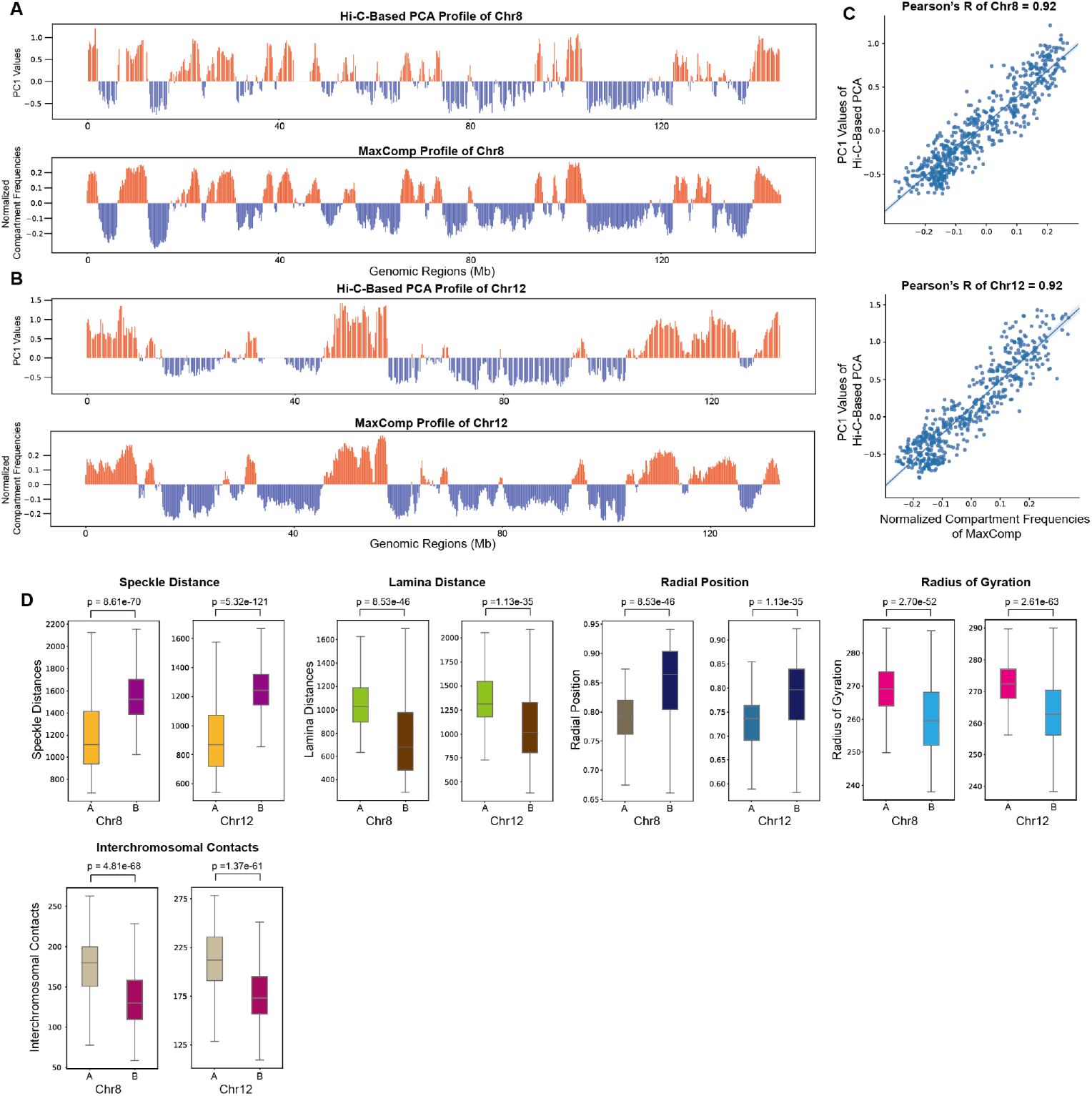
Prediction of model compartments by MaxComp and its comparison with the ground truth on H1-hESC Chr8 and Chr12. **A**, The experimental profile obtained from the Hi-C-based principal component analysis and the compartment profile predicted by MaxComp of 500 modeled structures of Chr8. **B**, The experimental profile obtained from the Hi-C-based principal component analysis and the compartment profile predicted by MaxComp of 500 modeled structures of Chr12. **C**, Scatter plot between the normalized compartment frequencies of MaxComp and the PC1 values together with the Pearson’s correlation coefficient between the two samples on each chromosome. **D**, Comparison of speckle distances (p-value=8.61e-70 and 5.32e-121), lamina distances (p-value=8.53e-46 and 1.13e-35), radial position (p-value=8.53e-46 and 1.13e-35), radius of gyration (p-value=2.70e-52 and 2.61e-63) and interchromosomal contacts (p-value=4.81e-68 and 1.37e-61) between compartment **A** beads and compartment **B** beads on the population of structures of Chr8 and Chr12.

**Figure S5:**
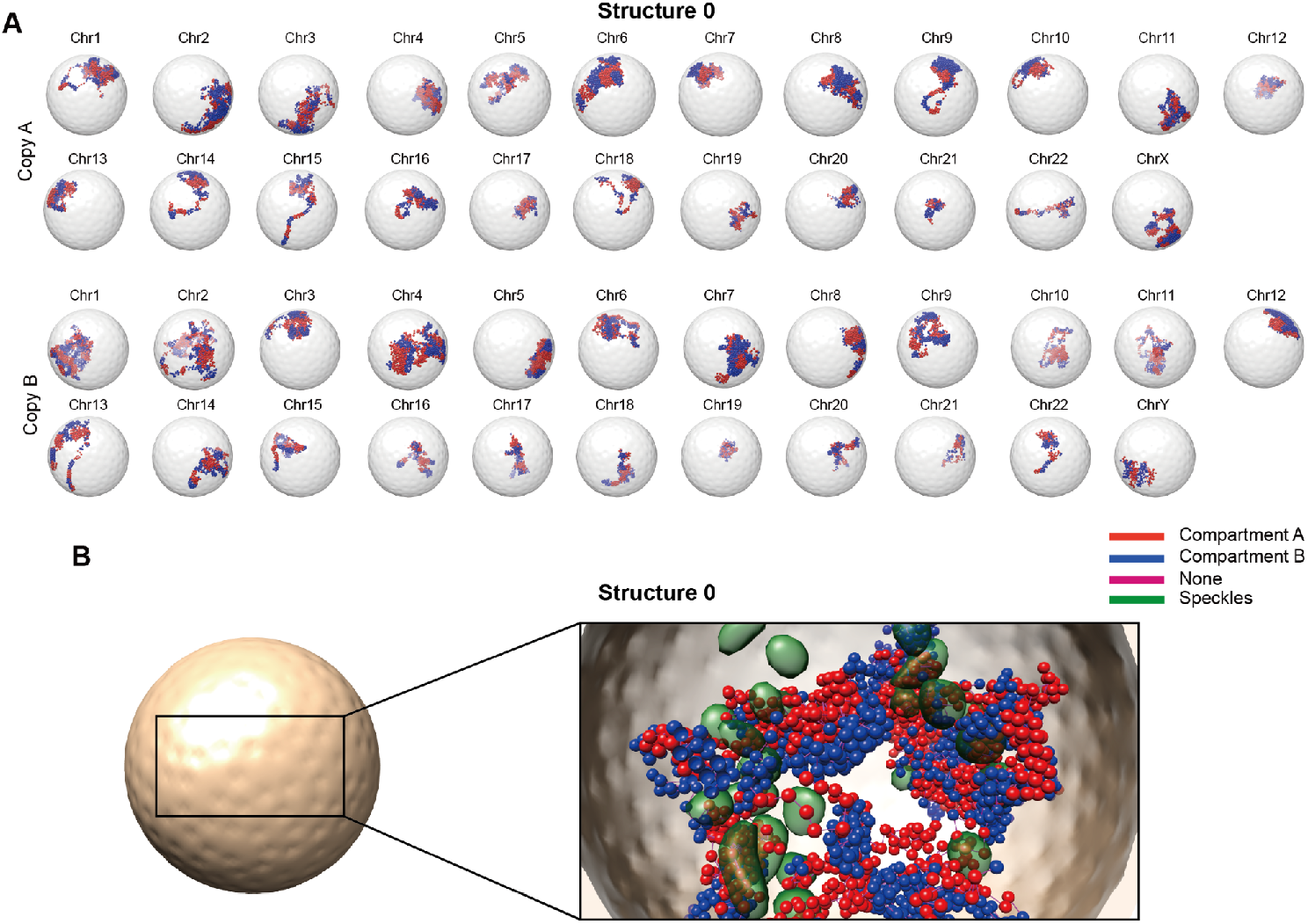
Selected example of model compartments predicted by MaxComp for the whole genome of H1-hESC. **A**, The predicted compartments for each chromosome copy from structure 0 of H1-hESC **B**, Selected chromosomes from structure 0 showed together with the envelope indicates compartment **A** and compartment **B** are segregated within the envelope. The section through the genome shows compartment **A** and compartment **B** are clustered with each other in the inferior region. Predicted speckles are basically associated with compartment **A** beads rather than compartment **B** beads.

**Figure S6:**
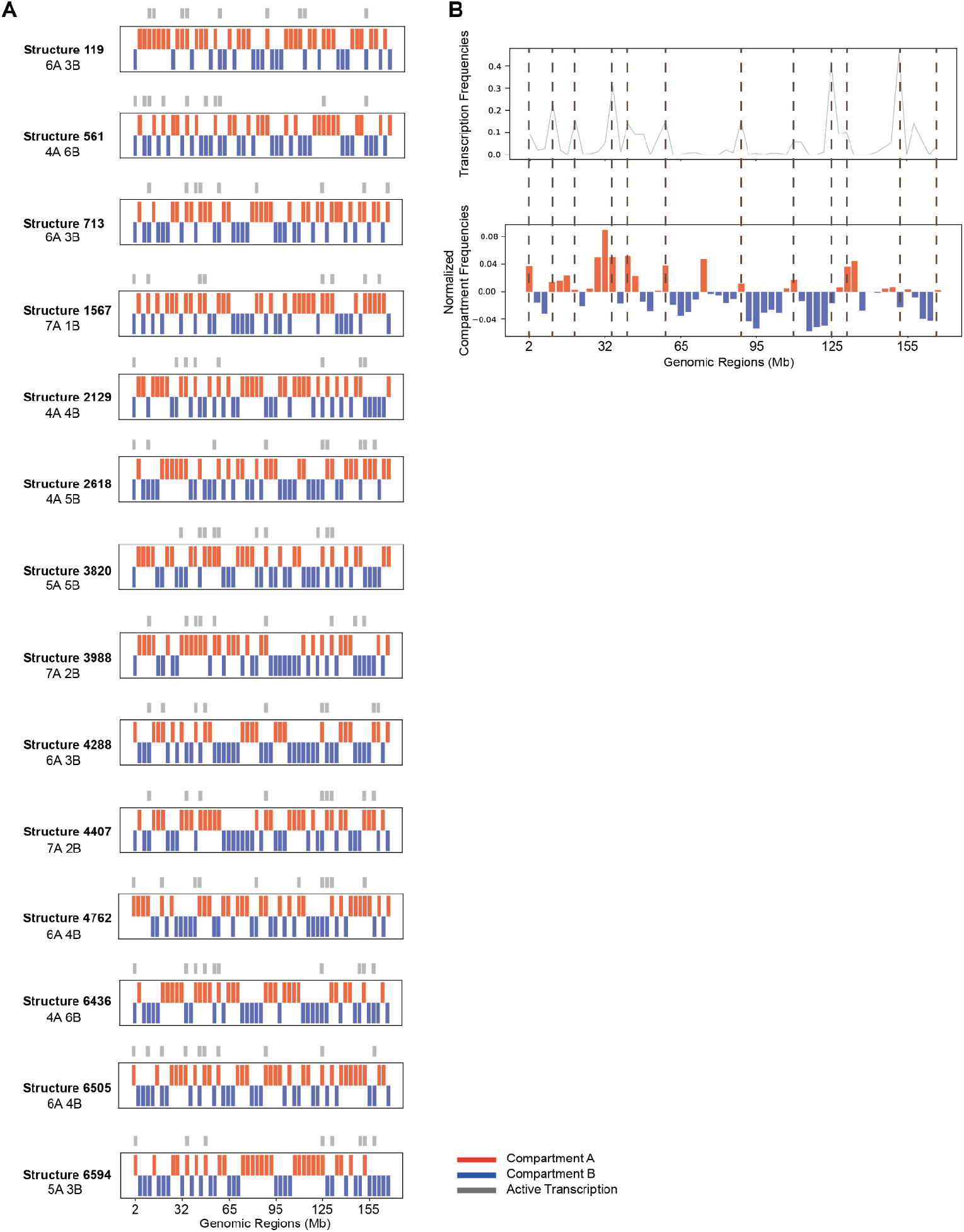
Transcription analysis on single-cell DNA-MERFISH Chr6 compartments predicted by MaxComp. **A**, Selected examples with more than or equal to 8 locus with active transcriptions of compartment prediction and transcription signals on DNA MERFISH structures [7] (Red bars indicate compartment **A**, blue bars represent compartment **B** while gray bars are where transcription is on (nascent transcript is imaged)). **B**, Comparison between the transcription frequency from DNA MERFISH and the compartment profile predicted by MaxComp.

**Figure S7:**
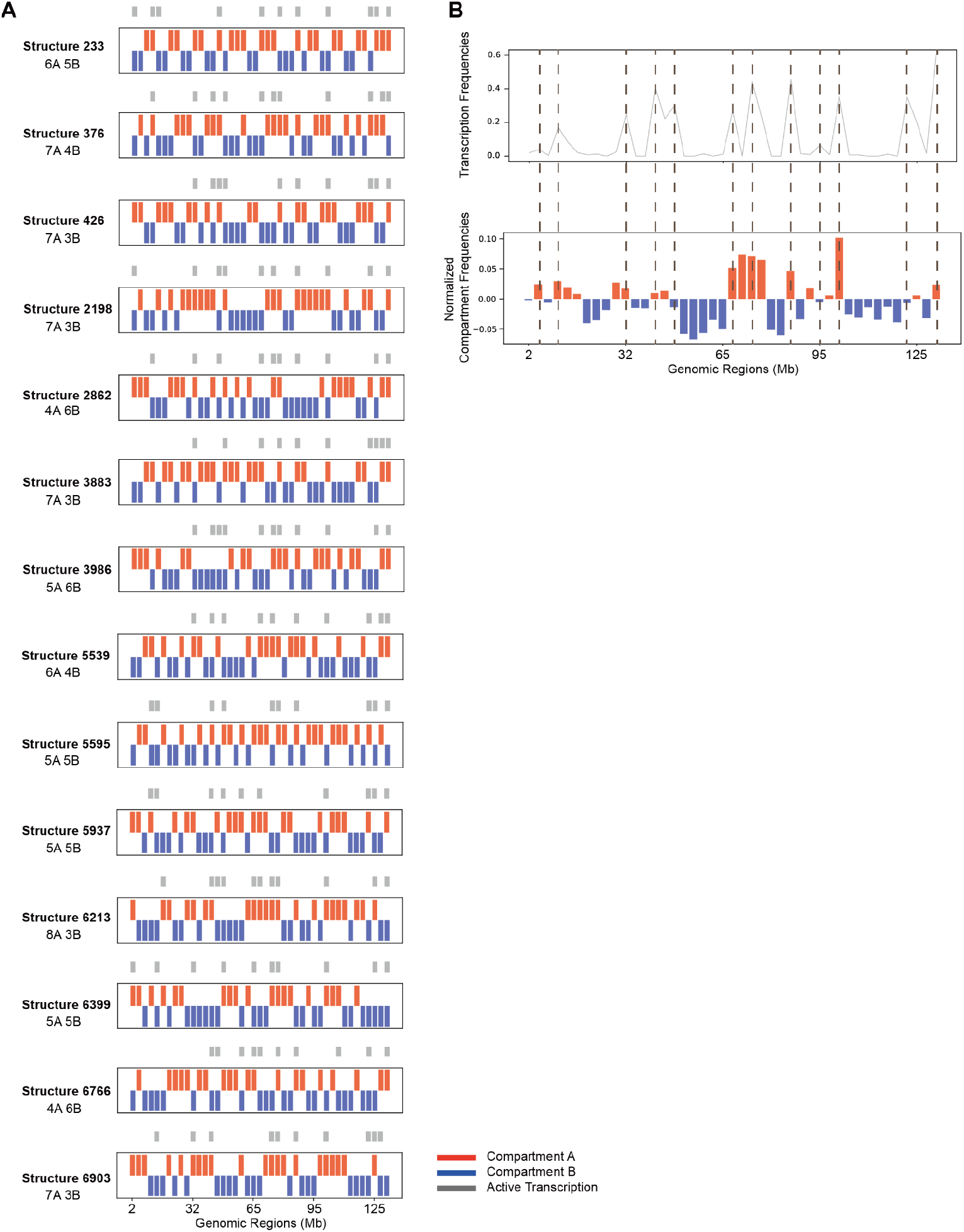
Transcription analysis on single-cell DNA-MERFISH Chr10 compartments predicted by MaxComp. **A**, Selected examples with more than or equal to 10 locus with active transcriptions of compartment prediction and transcription signals on DNA MERFISH structures [7] (Red bars indicate compartment **A**, blue bars represent compartment **B** while gray bars are where transcription is on (nascent transcript is imaged)). **B**, Comparison between the transcription frequency from DNA MERFISH and the compartment profile predicted by MaxComp.

**Figure S8:**
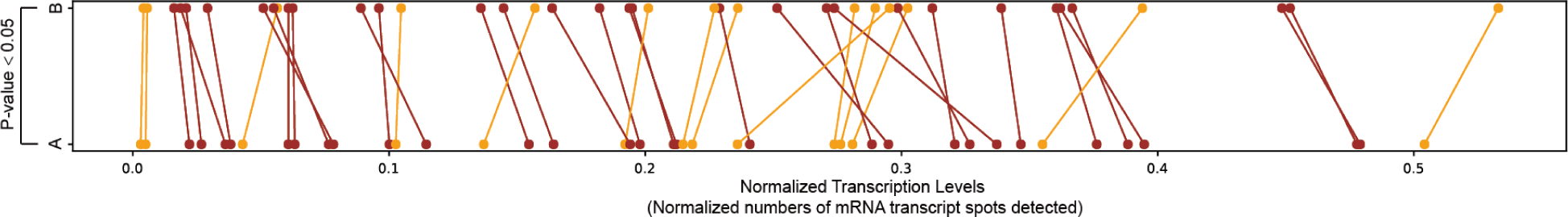
Comparison between distributions of normalized transcription levels from SeqFISH+. Comparison of normalized transcription levels (numbers of mRNA transcript spots detected) between **A** cells and **B** cells for genes measured by SeqFISH+ [8] showed together with the paired t-test. We find most of the genes have increased transcription levels when shifting from state **B** to state **A** (showed in brown).

